# Eco-evolutionary feedbacks and the maintenance of metacommunity diversity in a changing environment

**DOI:** 10.1101/2020.06.11.145680

**Authors:** Aidan P. Fielding, Jelena H. Pantel

**Affiliations:** Department of Biology, The College of William and Mary, P.O. Box 8795, Williamsburg, Virginia VA 23187-8795 USA; Department of Computer Science, Mathematics, and Environmental Science, The American University of Paris, 6 rue du Colonel Combes, 75007 Paris, France

**Keywords:** eco-evolutionary dynamics, eco-evolutionary feedback, metacommunity, evolving metacommunity, coexistence, competition, adaptive dynamics, quantitative trait evolution

## Abstract

The presence and strength of resource competition can influence how organisms adaptively respond to environmental change. Selection may thus reflect a balance between two forces, adaptation to an environmental optimum and evolution to avoid strong competition. While this phenomenon has previously been explored for single communities, its implications for eco-evolutionary dynamics at the metacommunity scale are unknown. We developed a simulation model for the evolution of a quantitative trait that influences both an organism’s carrying capacity and its intra- and interspecific competitive ability. In the model, multiple species inhabit a variable three-patch landscape, and we varied the connectivity level of the species among patches, the presence and pace of directional environmental change, and the strength of competition between the species. Our results reflect some patterns previously observed in evolving metacommunity models, such as species sorting and community monopolization. However species sorting was more likely to occur in evolving communities without dispersal, and monopolization was observed only when environmental change was very rapid. We also detected an eco-evolutionary feedback loop between local phenotypic evolution at one site and competition at another site, which maintains species diversity in some conditions. The existence of a feedback loop maintained by dispersal indicates that eco-evolutionary dynamics in communities operate at a landscape scale.

## Introduction

Competition for resources is a critical component structuring biodiversity in nature (Tilman 1994; Hautier et al. 2009). Many ecological theories developed to understand competition, coexistence, and biodiversity treat the traits that influence species interactions as fixed (e.g. Chesson 2000). Recent experimental studies have demonstrated, however, that critical parameters - such as the interaction strength of competitors (α) and the minimum resource level required for positive growth (R*) - can evolve on an ecological timescale (Hart et al. 2019; Bernhardt et al. 2020). Studies have also demonstrated that ecological competition can either inhibit or enhance evolutionary diversification and speciation, and can play an important role for understanding biodiversity over macroevolutionary time scales (Gillespie 2004; Losos & Ricklefs 2009; Drury et al. 2016). It is therefore important that theory is developed to predict how coexistence and consequent biodiversity are affected when traits influencing ecological competition can evolve (Roughgarden 1972; Lankau 2011). This is especially important as anthropogenic climate change increases existing or introduces novel selection pressures (Moran & Alexander 2014).

Theoretical models that evaluate the role of evolution for competition and coexistence in multiple species have been developed at both the local (single site) and regional (multisite) scale. At the regional scale, de Mazancourt et al. (2008) modeled phenotypic evolution in a trait that influences growth (where biomass at a site depends on the similarity between the population’s trait value and the local environmental optimum value), for multiple species that inhabited a patchy landscape and compete via a lottery for recruitment sites in proportion to their biomass. They found that adaptive evolution was slowed down when sites were connected, as local species were prevented from adapting by the arrival of pre-adapted immigrants (the metacommunity process of species sorting; Chase & Leibold 2003; Leibold et al. 2004). Other multisite studies used a similar approach, where the evolving trait determines fitness in the local environment and species compete via a micro-site lottery, and found that resident species can sometimes prevent colonization by other pre-adapted species due to a combination of local adaptation and a numerical advantage (community monopolization: Urban et al. 2008; Loeuille & Leibold 2008; Urban & De Meester 2009; Vanoverbeke et al. 2016). The study of Norberg et al. (2012) modeled competition differently, as per capita impacts of species on one another, and considered evolution in a trait that influenced growth rates (and was thus decoupled from competition). They found that local adaptation was inhibited by increased dispersal because species shifted their ranges instead of adapting to environmental change (which actually increased extinctions of species at range and trait extremes in their system). Taken together, these studies indicate that metacommunity composition emerges as a product of environmental heterogeneity, site connectivity (or species dispersal rate), and the speed of adaptive evolution. However, in all of these studies the measure of fitness (recruitment, survival, or intrinsic rate of population increase) depended only on the local environment and was decoupled from competition.

In nature, trait evolution can be influenced by both abiotic and biotic selection pressures. In zooplankton, for example, evolution of important life history traits such as body size is driven not only by temperature (e.g. Mc Kee & Ebert 1996), but also by selection for increased grazing efficiency (and thus increased competitive ability) and by predation risk from visually hunting fish (Burns 1969; Dodson 1974; Hall et al. 1976). Traits in protozoans, such as cell size or population growth rate, can also be influenced by competition and predation (terHorst et al. 2010; terHorst 2011). There are some models in which trait evolution is influenced by both the abiotic environment and ecological competition, but these consider only single sites. Nevertheless, these models have produced important findings that may have implications at the regional scale. In the adaptive dynamics model of Johansson (2008), the evolving trait influences both fitness in the local environment (i.e. carrying capacity depends on the distance between the trait value and the local optimum trait value) and the per capita effect of other species (i.e. the impact of competition depends on the difference in trait values between species). Competition was found to slow the rate of evolution, as trait convergence on an adaptive optimum determined by the environment led to increased competition with other species (and populations of the same species with differing trait values; Johansson 2008). However, the model of Osmond & de Mazancourt (2013) determined that competition can actually increase the rate of evolution in some instances - the degree of niche overlap between competitors (i.e. whether competition selects in the same direction as the new environment or in the opposite direction) determines whether competition slows or accelerates evolution. The goal of our study was to consider the effects of ecological competition on trait evolution in response to a variable environment at the regional, metacommunity scale. The combination of environmental heterogeneity and spatial connectivity could produce local communities that vary in their degree of niche overlap. When these communities experience a changing environment, their ability to adapt would thus be either diminished or accelerated. Spatial structure could thus play a critical role in the ability of competing species to avoid extinction and adapt to environmental change (evolutionary rescue; Gomulkiewicz & Holt 1995; Bell 2017).

We modeled population growth and competition for multiple species that inhabit a three-patch landscape. Population growth depends on a quantitative trait that determines both carrying capacity in the local environment and the degree of intra- and interspecific competition. This trait can evolve (implemented using a simulation of an adaptive dynamics framework; Metz et al. 1992; Dieckmann & Law 1996), and individuals can disperse among patches (Figure 1a). We monitored metacommunity eco-evolutionary dynamics to assess the effects of varying three components of the system: (1) spatial structure (isolated patches and patches with low and high levels of dispersal connectivity), (2) rate of environmental change (none, slow, and fast), and (3) strength of competition (by varying the decay in competition strength with increasing distance between individual trait values; Figure 1b). For each scenario, we evaluated evolutionary and community dynamics by considering local and regional species diversity, measures of stability, and distributions of the quantitative trait over time and across sites. Because the existing models of evolution and competition at the local and regional scale address some aspects of these components, we focused specifically on two questions. First, we evaluated whether niche overlap was a fixed property at each site or whether dispersal and environmental change caused this to vary over space and time. Niche overlap refers to the overlap between the niche of other competitors and the niche that a focal population is attempting to adapt to (Osmond & de Mazancourt 2013). When the direction of selection by competition and by the environment is completely overlapping – i.e. when the focal population is attempting to adapt to a niche already occupied by a competitor – adaptive evolution is constrained, but when niches partially overlap and competition decreases as the focal population adapts, the speed of adaptive evolution is actually increased. Osmond & de Mazancourt (2008) considered how fixed degrees of niche overlap in communities at a single location influenced the speed of evolution in their model, while Johansson (2008) varied the direction of environmental selection over time but not the degree of niche overlap. We were interested in using our metacommunity simulations to assess the existence and consequences of spatial and temporal variation in niche overlap. Second, we evaluated whether the findings of existing studies of evolution in metacommunities differed when trait evolution was influenced both by the environment and by competition. We used our evolving metacommunity model to determine whether the match between species traits and their environment (species sorting; Chase & Leibold 2003) observed at low to intermediate dispersal in previous studies is also observed when competition potentially restricts species from adapting to the local environmental optimum, and under what conditions global monopolization emerged.

**Figure 1.**
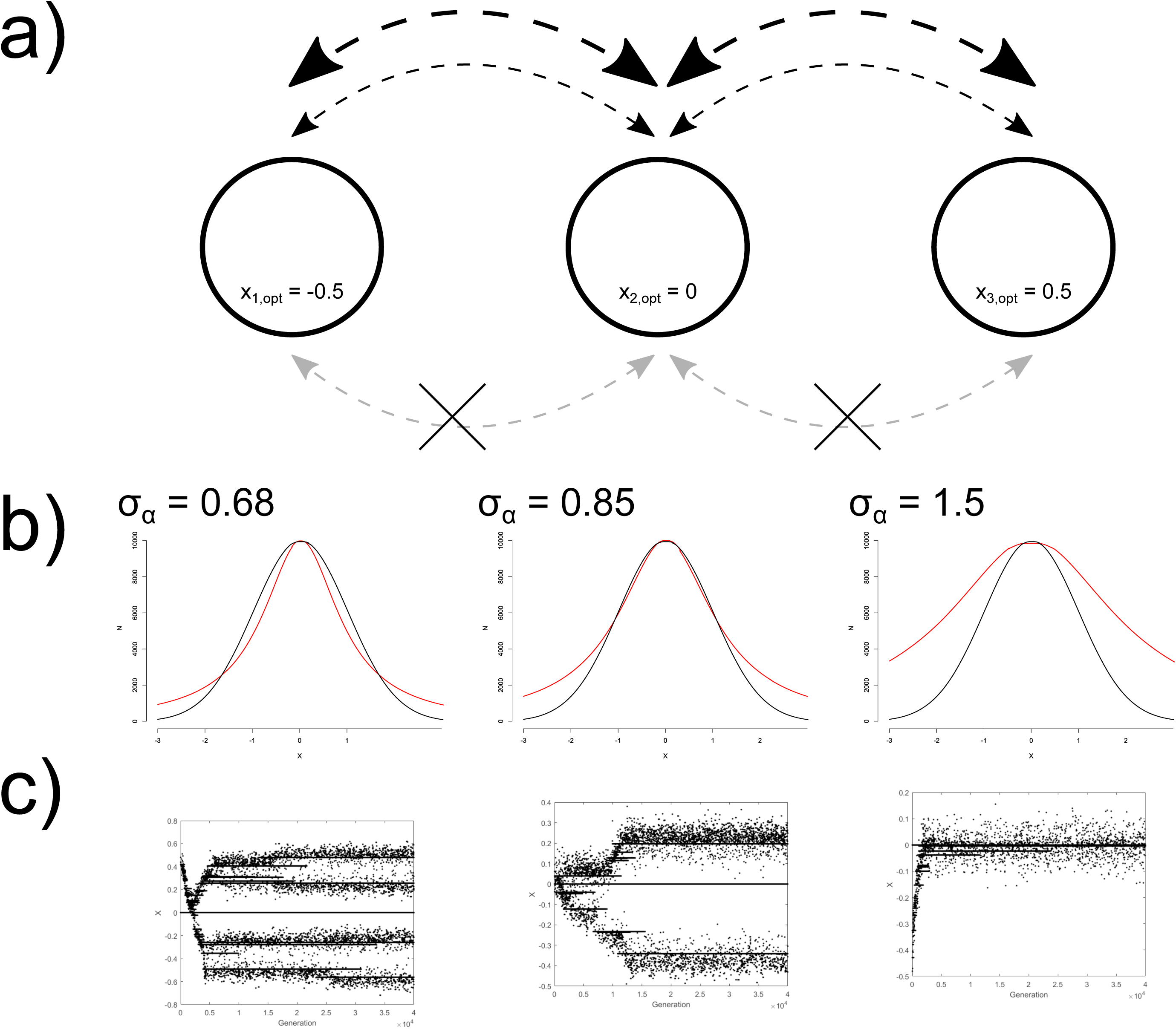
Model of metacommunity eco-evolutionary dynamics for species that compete for resources in a changing environment. (a) Communities inhabit a three-patch landscape with sites that vary in their resource distribution (which yields a distribution of carrying capacities centered around a maximum value with an optimal trait value *x*_*opt*_) and are connected at three levels of dispersal (*d* = 0, 0.01, 0.1 are the grey, thin black, and thick black lines respectively). (b) Simulations in the metacommunity were conducted using three different values for competition strength (σ_α_ = 0.68, 0.85, 1.5). Resident communities for the simulations were generated from modeling long-term coevolution in each patch (no dispersal) in a constant environment. The communities that emerge depend on the relationships between two functions that depend on a phenotype *x*. The competition function *C*(*x*) (red line) depends on the difference in trait values between two populations, with a standard deviation σ_α_. The carrying capacity function *K*(*x*) (black line) depends on the distance of a population’s trait from the local environmental optimum, can be interpreted as different resources that species can utilize at various efficiencies, and is a distribution with a standard deviation σ_K_ (equal to 1 in simulations). (c) The evolutionary history of the initial communities used in the simulation are shown. Species can coexist in a stable competitive hierarchy that evolves from a single initial population using a stochastic simulation of adaptive dynamics. In the plot, points represent trait values for all populations present at generation *t* (i.e. at least one individual exists with that trait value, population size is not represented), and a horizontal line at the optimal trait value is shown. The number of species (all unique populations in existence after 10^6^ generations) is determined by the ratio of the niche width to the width of the resource distribution (σ_α_ / σ_K_). An initial mutant population can evolve into multiple stable populations when σ_α_ / σ_K_ < 1 (corresponding to σ_α_ = 0.68 and 0.85 in this model), as a region of trait space exists where new mutants with trait values that differ from the local optimum still have a carrying capacity that exceeds the impact of competition (where the black line is above the red line). When σ_α_ / σ_K_ > 1 (σ_α_ = 1.5 in this model), a single population is maintained with a trait value at the local optimum.

## Methods

### Overview

We developed a model of quantitative trait evolution in a metacommunity of species that compete for resources and inhabit a three-patch landscape. Patches vary in their environmental properties (which favor an optimum trait value for resident species; Figure 1). Species in the model are defined as (asexually reproducing) populations with distinct phenotypes that vary along a single trait axis. Although it is important to consider multi-trait evolution, we first consider a single trait axis to expand from the results of existing models of evolution in competing species at the local and regional scale. We also note that the results are limited to understanding evolution in asexual populations with mutations of small effect (for a discussion of why the assumption of mutations with small effects may not accurately describe phenotypic evolution, and how sexual reproduction can influence competition for continuous resources, see Barton & Polechová 2005). Our model results will thus be most applicable to systems where a guild of asexually reproducing species respond to environmental selection pressures and compete for resources via a shared key trait such as body or beak size (e.g. Goulden et al. 1982; Lamichhaney et al. 2016), or resource uptake rate (e.g. Huston & DeAngelis 1994). Population growth follows Lotka-Volterra dynamics with density dependence and intra- and interspecific competition, and evolutionary dynamics are implemented using a simulation of the adaptive dynamics framework. This framework is useful to study effects of interactions between species that are linked by dispersal across multiple sites, because it links population dynamics directly to evolutionary dynamics (Dieckmann & Law 1996; Brännström et al. 2013), and also because these models can produce trait diversification that resembles distributions of species along niche axes observed in natural populations (Dieckmann & Doebeli 1999; Doebeli & Dieckmann 2000). The phenotypic trait determines an individual’s carrying capacity and the strength of competition with other individuals. We model environmental change as a constant rate of increase in the optimum trait value (the trait value that leads to the highest carrying capacity). We consider the same rate of change for all patches. All individuals of all species have a uniform probability of dispersal and can move equiprobably to the other patches in the landscape. The formulas and model details for ecology, evolution, and dispersal are described below in the sections *Within-patch ecology, Evolution, Extinction, Dispersal*, and *Environmental dynamics*. The details for simulation conditions to address our research questions are described in the sections *Metacommunity initialization* and *Simulation conditions*. Finally, we describe measures for quantifying eco-evolutionary properties (e.g. diversity, degree of phenotypic evolution) in the section *Metrics for evolutionary and ecological properties*.

### Within-Patch Ecology

Population dynamics are evaluated for all populations *i* that differ in their trait value *x*, i.e. each population has its own trait value *x*_*i*_ and population size *N*_*it*_ varies over time *t*. Novel phenotypes that arise via mutation are considered a new population. Population size *N*_*it*_ is a function of the vectors of trait values **x** for all populations present in the local patch and their population size in the previous generation *N*_*jt-1*_. Population growth depends on trait values **x** in two ways. First, the carrying capacity *K* of a population with trait value *x*_*i*_ is a Gaussian function of the distance of *x*_*i*_ from *x*_*opt*_, which is the patch-specific optimal trait value:

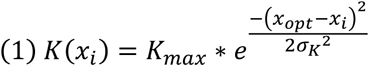

Here *x*_*opt*_ is the trait value corresponding to the maximum carrying capacity, *K*_*max*_, and *K*_*max*_ is the same for all populations in the patch. Carrying capacity decreases if population traits are distant from the optimum *x*_*opt*_, and the width of this carrying capacity distribution is determined by the standard deviation term σ_*k*_. *K*(*x*) can be interpreted as representing a resource distribution (biotic or abiotic) on a given patch, scaled to equal the total population that the resources at any given position on the trait axis can sustain. A population’s trait *x*_*i*_ therefore corresponds to its ability to exploit its most efficiently used resource, and its carrying capacity *K*(*x*_*i*_) is limited by that resource.

Competition between populations is a symmetric function of the difference in their trait values. Competition, the per capita negative effect of a population with trait *x*_*j*_ on a population with trait *x*_*i*_ (α_*ij*_) is determined by a decaying symmetric function of the distance between trait values *x*_*i*_ - *x*_*j*_:

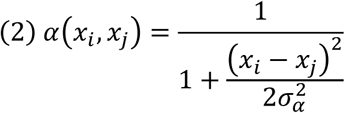

where the strength of competition between individuals with differences in trait values is determined by the standard deviation term σ_*α*_ (i.e. the niche width; we subsequently refer to the value of σ_*α*_ as the competition strength or as the niche width). When *x*_*i*_ = *x*_*j*_ the populations are ecologically equivalent and their competitive effect on each other is the same as their intraspecific effect on themselves (α_*ii*_ = 1; otherwise 0 ≤ α_*ij*_ < 1).

Equations 1 and 2 are incorporated into each population *i*’s growth:

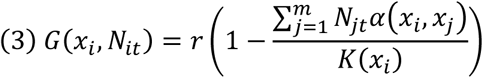

Where *r* is the population’s intrinsic rate of increase and *m* is the total number of populations present in the patch, across all species (more on how species are defined in the section *Metacommunity initialization*). We considered discrete population dynamics, so population size from one time step to the next is:

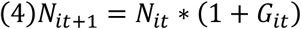

### Evolution

A mutation rate µ gives the probability that an individual born in a population carries a mutation. In our model, existing populations give birth to *rN*_*it*_ individuals every generation. Each mutant born in a population in a given generation (total number of mutants *N*_*m*_ = µ*rN*_*it*_) is treated as a new population (subtracted from the parent population size, with initial population size equal to 1), and mutants do not encounter competition in the generation they are generated (which allowed for computational tractability of tracking all mutant populations over time; results with competition in the same generation did not qualitatively differ from those presented here). For all populations, mutants are subtracted from all individuals born to a parental population (*rN*_*it*_) before the non-mutant individuals in the parental population experience competition (i.e. before the terms in the parentheses in Equation 3 are applied). The trait value of the mutant population, *x*_*m*_, is drawn from a normal distribution centered at the parental trait value *x*_*i*_ with standard deviation σ_µ_, 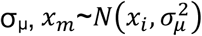.

### Extinction

At the end of each generation *t*, we remove all populations with *N*_*it*_ < 1. We also implemented stochastic extinction at low population size: all populations with *N*_*it*_ below an extinction threshold θ but above 1 have a probability ρ of being removed from the system.

### Dispersal

All patches are equally connected by dispersal. Each generation, all individuals can disperse with probability *d*. Dispersing individuals from one patch have an equal probability of migrating to any of the other patches.

### Environmental dynamics

The metacommunity landscape consists of *k* patches (in all of the simulation conditions considered in this study, we set k = 3). The optimal trait values *x*_*opt,k*_ are evenly spaced along the trait axis with fixed difference δ, which gives the degree of environmental heterogeneity in the metacommunity. Environmental change was implemented each generation, where the patch optimal trait values *x*_*opt,k*_ increase by the same rate Δ in all patches.

### Metacommunity initialization

We used equations 1-4 to simulate adaptive dynamics in a single initial population with *N*_0_ = 500, *x*_0_ = 0, and *K*_*max*_ = 10000. This initial population was placed in each of *k* = 3 patches (which represent the patches used subsequently in the evolving metacommunity simulation; Figure 1a), using among-patch environmental distance δ = 0.5. Local environmental optima in each patch *k* (*x*_*k,opt*_) were therefore *x*_1,*opt*_ = −0.5, *x*_2,*opt*_ = 0, and *x*_3,*opt*_ = 0.5. For this initialization, we also set *d* = Δ = 0, σ_*k*_ = 1, *r* = 1.9, µ = 10^-5^, and σ_µ_ = 0.05, θ = 2, and ρ = 0.025 (results were qualitatively similar but over longer time scales for lower values of µ and σ_µ_. We generated three distinct initial metacommunities, using each of three levels of niche width (σ_α_ = 0.68, 0.86, 1.5). Evolutionary dynamics proceeded for 10^6^ generations, to create a community of species that have co-evolved to be well suited for their local environment and for coexistence in competition with the other species (Figure 1c). The adaptive dynamics framework provides a mechanism for emergence of multiple distinct lineages from a single initial trait value (described in more detail in Results, *Initial communities*). The coexisting lineages present at the end of the initialization period were considered unique species (similar to Johansson 2008) and are stochastic approximations of the coevolutionary equilibrium for that set of model conditions.

### Simulation conditions

Coevolved communities that evolved from initial conditions in the three patches independently for 10^6^ generations were used as initial patch communities for the evolving metacommunity simulations. For each of the three different levels of niche width (σ_α_), we ran simulations for each three-patch metacommunity at different dispersal levels (*d* = 0, 0.01, 0.1) and different rates of environmental change (Δ = 0, 10^-5^, 4×10^-4^), maintaining the same level of niche width for that metacommunity. All of the 27 simulations ran for 50,000 generations and all simulation conditions were introduced at *t*_0_. All simulations were conducted in MATLAB R2017a version 9.2.0.556344.

### Metrics for evolutionary and ecological properties

A species *l* is defined as a population with a unique trait value at *t*_0_ in the simulation. All populations *m* that belong to a species (i.e. that arose via mutation from the initial population) are considered sub-populations of that species (and so individuals with particular trait values can be indexed as *x*_*lm*_). Each generation, we calculated patch alpha diversity using the inverse of Simpson’s diversity index: 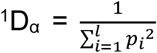 where *l* is the number of species present in site *k* and *p*_*i*_ is the relative proportion of the species at that site (all populations *m* for a species *l* are included in this *p*_*i*_ calculation). We calculated regional gamma diversity ^1^D_γ_ using the same formula, but after summing population sizes of each species across all sites, and we calculated beta diversity as 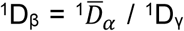. We calculated the coefficient of variation in total population size within a patch (i.e. summed across all populations for all species, giving a CV for the total number of individuals in a patch) and the sum of squared deviations of individual trait values from the local environmental optimum (calculated for all individuals *n* within a patch; 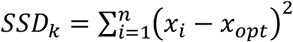

To better understand how equations 1 and 2 interacted to structure evolutionary trajectories in our evolving metacommunity, we generated approximations of the functions of population loss due to competition *C*(*x*) and the carrying capacity *K*(*x*), depending on trait value *x*, for each generation (i.e. the curves seen in Figure 1b). The *C*(*x*) curve gives the distribution of intra- and interspecific competitive effects across the trait axis, 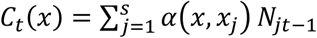, for all populations *s* at time *t* (calculated at discrete values of *x* in intervals of 0.001 for the range {*x*_*opt,k*_ - 3, *x*_*opt,k*_ + 3}). The value for an existing population’s trait value (i.e. a population actually present in the patch at that time) in generation *t* is calculated including both intraspecific competition (with other individuals with the same trait value) and interspecific competition (with individuals with all other trait values, across the same and other species) experienced in generation *t*, and for all other *x* values in the range it is calculated as the total amount of competition with individuals that have differing trait values. The *K*(*x*) curve was calculated using equation 1 for the range of *x* values. Populations with trait values in a region of the trait axis where *K*(*x*) > *C*(*x*) experience positive population growth in generation *t*, whereas those with *K*(*x*) < *C*(*x*) experience negative growth (equation 4). All metrics and figures were made using R version 3.5.0 (2018).

## Results

### Initial communities

Our model is a stochastic simulation of an adaptive dynamics model (Metz et al. 1992; Dieckmann & Law 1996), and previous studies have found evolutionary branching, where populations with multiple trait values can co-exist despite the presence of a single phenotypic optimum (Geritz et al. 1998; Kisdi 1999). Competition in our model is symmetric, (Christiansen & Loeschcke 1980; Johansson 2008), which means that the ratio between niche width and the width of the resource landscape (σ_α_ / σ_K_) determines when multiple populations can stably co-exist in the absence of dispersal (Figure 1b, c). Evolutionary dynamics and consequent species composition in our model is thus determined by the position of both symmetric functions *α*(*x*) and *K*(*x*) on the resource optimum *x*_*opt*_, and also the shape of the functional form for *α*(*x*).

We used the competition function given in equation 2 as described in Johansson (2008), which has heavier tails than the Gaussian curve used for *K*(*x*). We evaluated the impacts of competition for population size using the function of population loss due to competition *C*(*x*). When niche width is smaller than the width of the resource distribution (σ_α_ < σ_K_) and the resident population is adapted to the resource optimum (*x*_*i*_ = *x*_*opt*_), there are regions of trait space where *K*(*x*) exceeds *C*(*x*), and thus mutants with trait values away from the optimum can have positive population growth. In contrast, 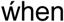 σ_α_ > σ_K_ no mutants can invade, as the reduction in competition is insufficient to compensate for the reduced carrying capacity (Dieckmann & Doebeli 1999; Doebeli & Dieckmann 2000; Figure 1b). In all cases, the evolutionarily stable configuration of species is characterized by a balance between two opposing selective forces: one for increased carrying capacity and one for decreased competition. As niche width decreases relative to the width of the resource landscape, the number of evolutionarily stable coexisting populations and the variance of those populations in trait space both increase. We varied niche width in our simulations using σ_α_ = 0.68, 0.85, and 1.5 (and σ_K_ = 1), which respectively produce 4, 2, and 1 distinct branches as evolutionarily stable solutions in trait space (Johansson 2008; Figure 1c). Initial combinations of species can be variable in some ways in different realizations of our stochastic simulation – the positions of species around the local *x*_*opt*_ varied when the initialization step was repeated (data not presented here), and the number of distinct branches were metastable, in that differing numbers of branches persisted for very long periods of time before shifts to branch numbers closer to the predicted evolutionarily stable number emerged. The number of branches became more repeatable the longer the initialization step was run, and all of our repetitions of the initialization step for 10^6^ generations had the same number of branches as those that were used in this study. Simulations to address our two main research questions subsequently used these communities in each patch for the *t*_0_ metacommunities. We thus refer to each distinct phenotypic value persisting at generation 10^6^ after the initialization as a unique species.

### Effect of spatial structure for trait evolution and metacommunity diversity

To determine how spatial structure influences metacommunity eco-evolutionary dynamics, we evaluated phenotypic distributions and species composition over time when dispersal *d* = 0, 0.01, and 0.1 for *t* = 50,000 generations. To evaluate diverse metacommunities where species can persist in multiple sites, we considered Δ = 0, σ_α_ = 0.68, and δ = 0.5 for all results unless otherwise stated (results to address the effects of varying Δ are discussed in section *Environmental change* below and of varying σ_α_ in the section *Increasing competition strength* below).

### 1. Shifts in direction of selection by environmental optimum and competition

Dispersal introduces individuals with distinct trait values (Figure 2), which alters the *C*(*x*) curve and the resulting selection pressures. Dispersal slightly increases phenotypic homogenization towards the overall metacommunity 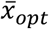 (i.e. small populations with higher trait values colonize and persist in patch 1 and with lower trait values in patch 3, although these immigrant populations are at low density; Figure 2a, d > 0) and increases local community phenotypic variance (the distance between the minimum and maximum trait value present in a patch; Figure 2a). Immigrants add population density to their region of trait space, which shifts the distribution of competitive effects *C*(*x*) towards the distribution’s tails (as persisting immigrants - those with positive growth in the new patch - in patch 1 tend to have higher trait values than residents, persisting immigrants in patch 2 tend to have both lower and higher trait values than residents, and persisting immigrants in patch 3 tend to have lower trait values than residents). This is most easily seen in Figure 2b at high dispersal (*d* = 0.1). The region of trait space in each patch where species can have positive realized growth rates (*x*_*fit*_, which we define as the region where *K* > *C*) can thus vary among patches, and the width of this region increases with dispersal. Populations with traits in these higher-fitness regions are selected to be maintained (i.e. populations with trait values in the region where *K* - *C* was highest had the highest population sizes; these values approximate the expected trait equilibrium indicated in Osmond and De Mazancourt 2013; Figure 2b). Dispersal shifts local phenotypic distributions in the direction of immigrant trait values, so patch 1 has a high fitness region for lower trait values and patch 3 has a high fitness region for higher trait values (i.e. in the direction where mutants experience less competition). Increasing dispersal increases the width (the range of trait values *x*_*fit*_) and depth (*K*(*x*_*fit*_) - *C*(*x*_*fit*_)) of the positive growth region. The direction of the competition curve in patch 2 does not shift (and thus resident populations with *x*_*opt*_ continue to experience the most competition), but its height in the center is reduced due to immigrants increasing the strength of competition in the right and left tails of the *C*(*x*) distribution.

**Figure 2.**
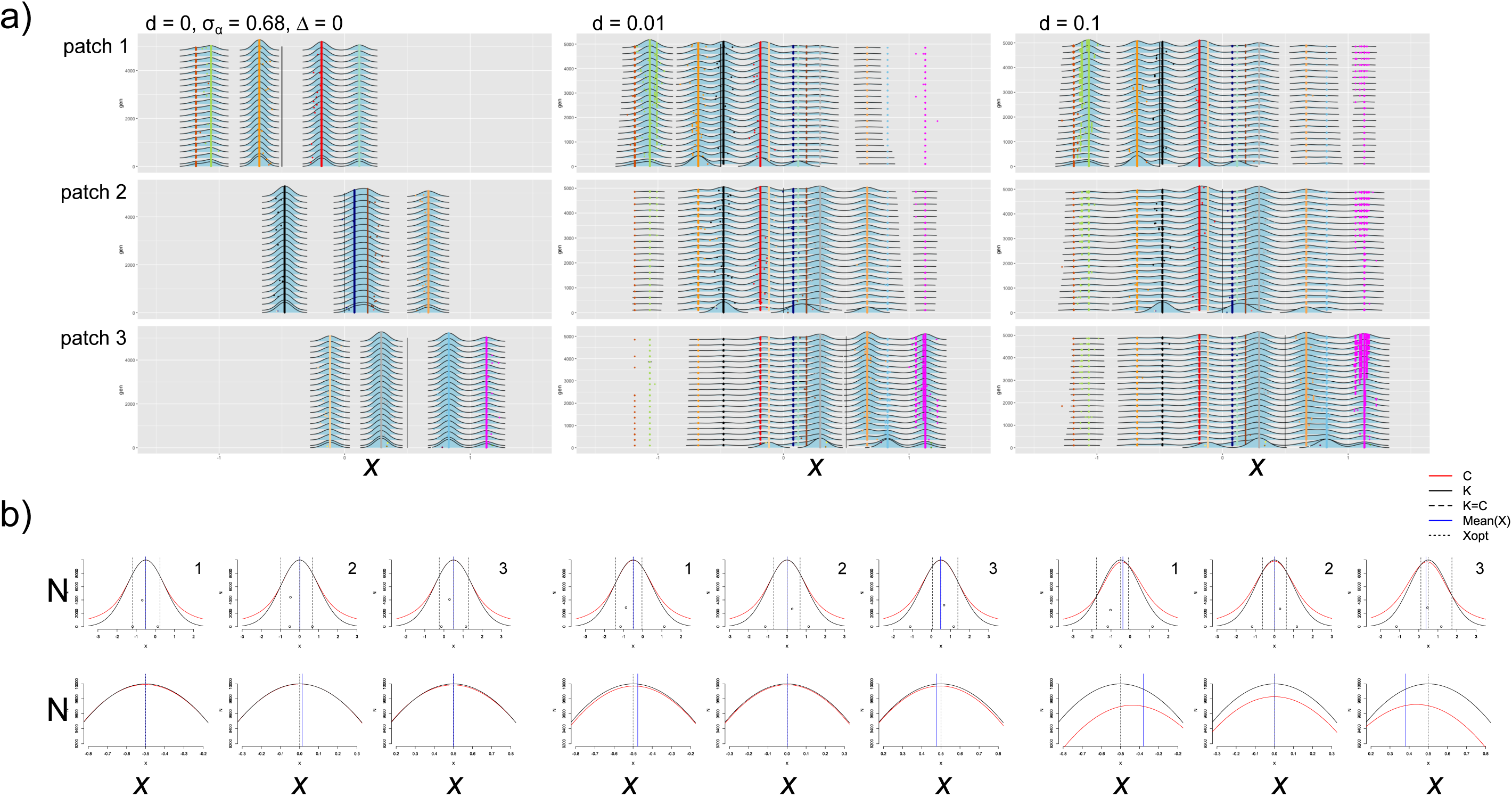
Trait evolution and competition for multiple species in communities over time in an unchanging environment (Δ = 0, σ_α_ = 0.68, *d* = 0, 0.01, and 0.1). (a) Time series of phenotypic distributions. Density plots of trait values (*x*-axis) weighted by relative abundance (height of density curve) are shown over 5,000 generations (*y*-axis; the time series was thinned for visualization, so density plots along the *y*-axis are shown every 250 generations), with a colored point indicating the species identity of lineages in each generation (each species is assigned a unique color). Simulations were run for 50,000 generations but the dynamics are clear within the first 5,000 generations. The solid black line in each plot indicates the local optimum. (b) Competition strength and carrying capacity as a function of trait value for *d* = 0, 0.01, and 0.1 (left to right) in three patches (left to right are patch 1, 2, and 3). Plots on the top row show competition (*C*(*x*), red) and carrying capacity (*K*(*x*), black) as a function of trait value *x* (*x*-axis) and are calculated as average values from 121 sampled generations spaced evenly from generation 20,000 to 50,000. Black points show the phenotypic distribution in each patch and their corresponding population size (*y*-axis). The left and right-most points are the average trait values (across the 121 generations considered from time 20,000 to 50,000) of the species with the lowest and highest trait values and the middle point is the average trait value of the species with the highest population size. The patch’s *x*_*opt*_ (short-dashed black line), the community’s mean *x* (blue solid line, averaged across the sampled 121 time points, weighted by population size), and points where the carrying capacity and impact of competition are equal to one another (long-dashed black line) are shown as well. The plots on the bottom row are zoomed in to better see distinctions between the *C*(*x*) and *K*(*x*) functions.

### 2. Phenotypic distributions, diversity, stability, and degree of maladaptation

Introducing novel species via dispersal led to species replacement and altered local community composition (Figure 2a). In each patch, at least one resident species was replaced by an immigrant species. In patch 1, when *d* = 0.01, an immigrant species from patch 2 with a trait very close to the local resource optimum (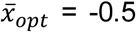, colored black) establishes. The presence of established immigrant species alters the local fitness landscape, and new combinations of species (including the successful establishment of a species from patch 3 colored light yellow, the eventual loss of a resident species in patch 1 colored red, and the strong reduction in population size of the patch 1 resident with the highest trait value colored light blue; extinctions can be observed in Figures 3, S5, and S6, which show phenotypic distributions for 50,000 generations). In patch 2, two resident species with traits closest to the local environmental optimum (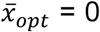, colored blue and brown) are either lost or greatly reduced in population size as immigrants from patch 1 (colored red) and patch 3 (colored grey) are favored in the novel fitness landscape introduced by dispersal. In patch 3, the species combinations change as two resident species (colored light orange and light blue) are replaced by immigrants from patch 1 (colored red) and patch 2 (colored orange). The precise trait values of the novel species combinations are difficult to predict, but they differ from the combinations observed to be metastable after the 10^6^ generations of the initialization step. This indicates that dispersal introduces a fitness landscape among competitors that differs from the one observed when sites are isolated (Figure 2b). With a higher colonization rate (*d* = 0.1), the fitness landscape shifts and there is a wider region of trait space where populations have positive growth (Figure 2b). As a result, some species that were lost to extinction at the lower dispersal level are now retained (e.g. the species colored red in patch 1, the species colored brown in patch 2, and the species colored light blue in patch 3; Figure 2, Figure 3, Figures S5 and S6).

**Figure 3.**
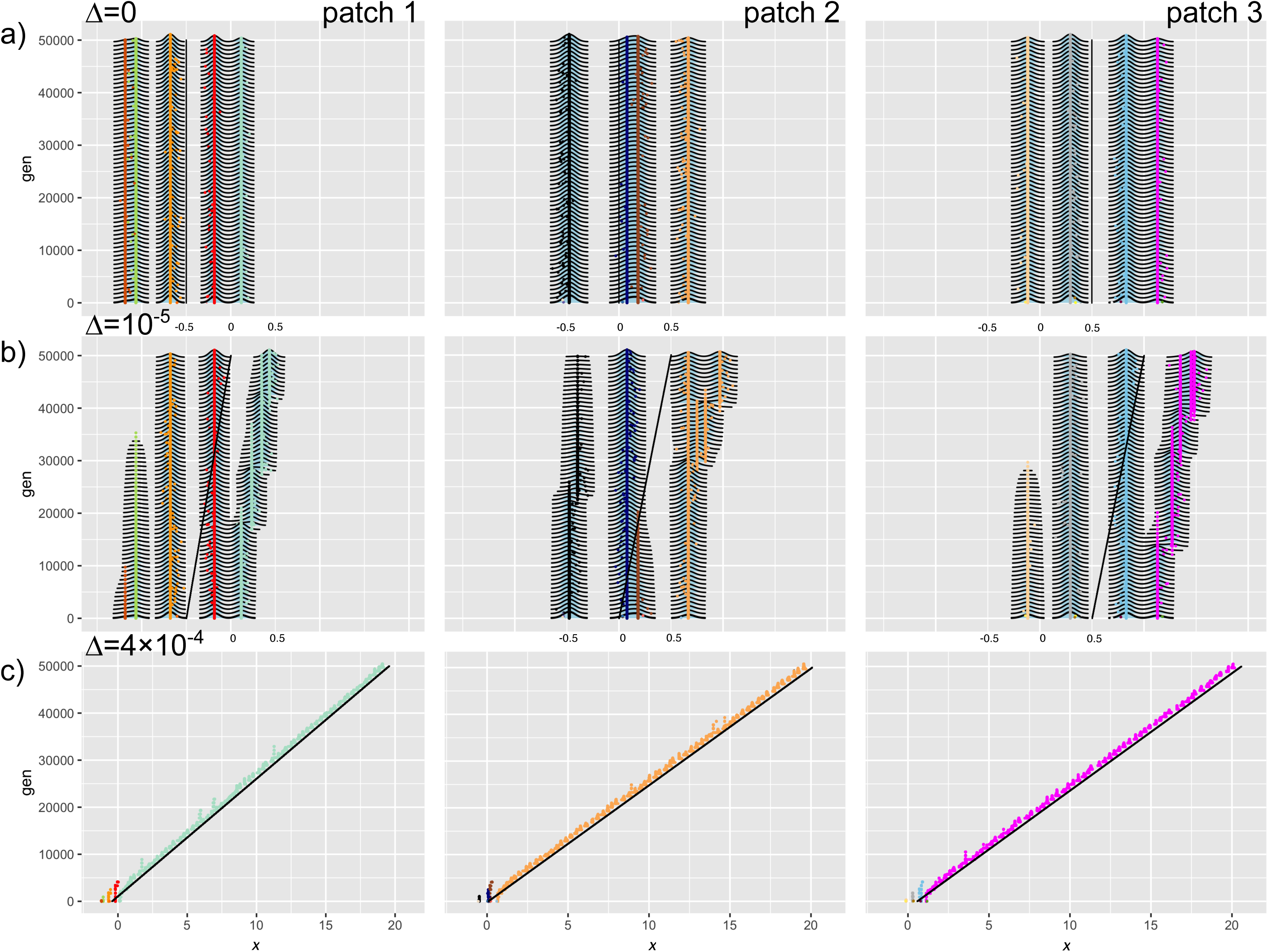
Trait distributions in communities over time across a range of environmental change rates. In all plots, σ_α_ = 0.68 and *d* = 0, and the rate of environmental change varies: (a) Δ = 0, (b) Δ = 10^-5^, and (c) Δ = 4×10^-4^. The *x*-axis range differs for (c), as traits span a wider range (see Figure S4 to better compare width of phenotypic distributions across all three rates of environmental change). Communities in patches 1, 2, 3 are shown (from left to right) and features of the trait density plots are as described in Figure 2a (except that the time series are thinned to show every 800 generations). In (b), the species with the highest trait value in each patch is less constrained by competition than other species and experiences the most adaptive trait change. Species with lower trait values are gradually lost to extinction. In (c), all species are lost within 5000 generations and the species with the initially highest trait values persists and adapts over time.

The loss of resident species with dispersal leads to a decrease in regional γ diversity, and replacements by immigrant species causes local α diversity to increase, converging to regional γ diversity (Supplementary Information Figure S1). Diversity is higher at *d* = 0. 1 compared to *d* = 0.01. As dispersal increases the width of the region of trait space with positive population size (Figure 2b), more mutants are maintained with increasing dispersal and so intraspecific diversity increases with dispersal (Figure S1). Some species with trait values very similar to other species (i.e. the species colored brown and light blue in Figure S5) were lost to extinction, which reduced the competition for mutants of other species (i.e. the species colored green and magenta in Figure S5). However in the absence of environmental change, where most species are pre-adapted to their environment, broad shifts in community trait composition are primarily due to species replacements and there are fewer shifts in a given species trait distribution. Dispersal leads to slight increases in SSD, as immigration pressure allows more species to persist in different patches, including those distant from the local trait optimum. The patches experience more shifts in population size with dispersal (and thus increasing CV), as the shifted fitness landscape leads to gradual reductions in some populations and increases in others (Figure S1).

### Environmental change

#### 1. Shifts in direction of selection by environmental optimum and competition

To understand the effects of environmental change, we first evaluated simulation results for *d* = 0 (no dispersal), σ_α_ = 0.68, and δ = 0.5 with varying levels of environmental change Δ = 0, 10^-5^, and 4.4×10^-4^. Directional environmental change (increased *x*_*opt*_ each generation) shifts the carrying capacity curve *K*(*x*) to the right at all sites, selecting for individuals with higher trait values (Figure S2). Over time, mutants with trait values higher than the rest of the community experience minimal competition, which increases the overall selection for increasingly higher trait values as environmental selection and selection for reduced competition both favor traits in the same direction. This partial niche overlap – where competition is decreased in the direction selected for by the environment – reflects the partial niche overlap described in Osmond & de Mazancourt (2013; see their Figure 6a), where the speed of evolution is actually enhanced by competition.

#### 2. Phenotypic distributions, diversity, stability, and degree of maladaptation

In our simulations, the addition of relatively slow environmental change (*d* = 0, Δ = 10^-5^) caused phenotypic distributions to have gradually increasing trait values over time (Figure 3a and b). Species with the lowest trait value in a patch persisted at an increasingly reduced carrying capacity until they were no longer viable in the changing environment and went extinct, as mutants in these populations with increased trait values experience intense competition from other species with similar trait values. Adaptation thus primarily occurred in the species with the highest trait value (Figure 3b, colored magenta, orange, and light green), as most of the other species in a patch continue to experience complete niche overlap (where competition is increased in the direction of selection). Gradual environmental change thus led to an overall decrease in species diversity due to extinctions, but an increase in intraspecific diversity (Figure S3). Gradual environmental change had no strong changes on SSD (compared to no environmental change), and the CV was increased when *d* = 0, likely due to the changing population sizes of increasingly maladapted species that are eventually lost to extinction and of increasingly adapted species as well (Figure S3).

With rapid environmental change (*d* = 0, Δ = 4.4×10^-4^) all species in each patch except that with the highest trait value went extinct relatively quickly (Figure 3c; Figure 4a). Though interspecific diversity is lost (Figure S8) due to the stronger selection pressure against species with lower trait values (Figure S2b), intraspecific diversity for the remaining species is higher than with slow environmental change (Figure S9; Figure S4). This is because selection for increased trait values and decreased competition combine to create a large region of positive carrying capacity (where *K* > *C*), and many mutants with very different trait values higher than residents can persist (Figure S2b). As was observed by Johansson (2008), the single remaining species in each patch can track the changing environment and the mean trait value lags behind the optimum (e.g. Figure 4a; this is similar to models without competition, e.g., Bürger & Lynch 1995; see Kopp & Matuszewski 2014 for a review of quantitative trait evolution in response to environmental change). As the rate of environmental change increases, the overall metacommunity SSD thus decreases and metacommunity CV increases (because the numerous surviving mutants vary in their population size as the environment continues to change; Figure S10 and S11).

**Figure 4.**
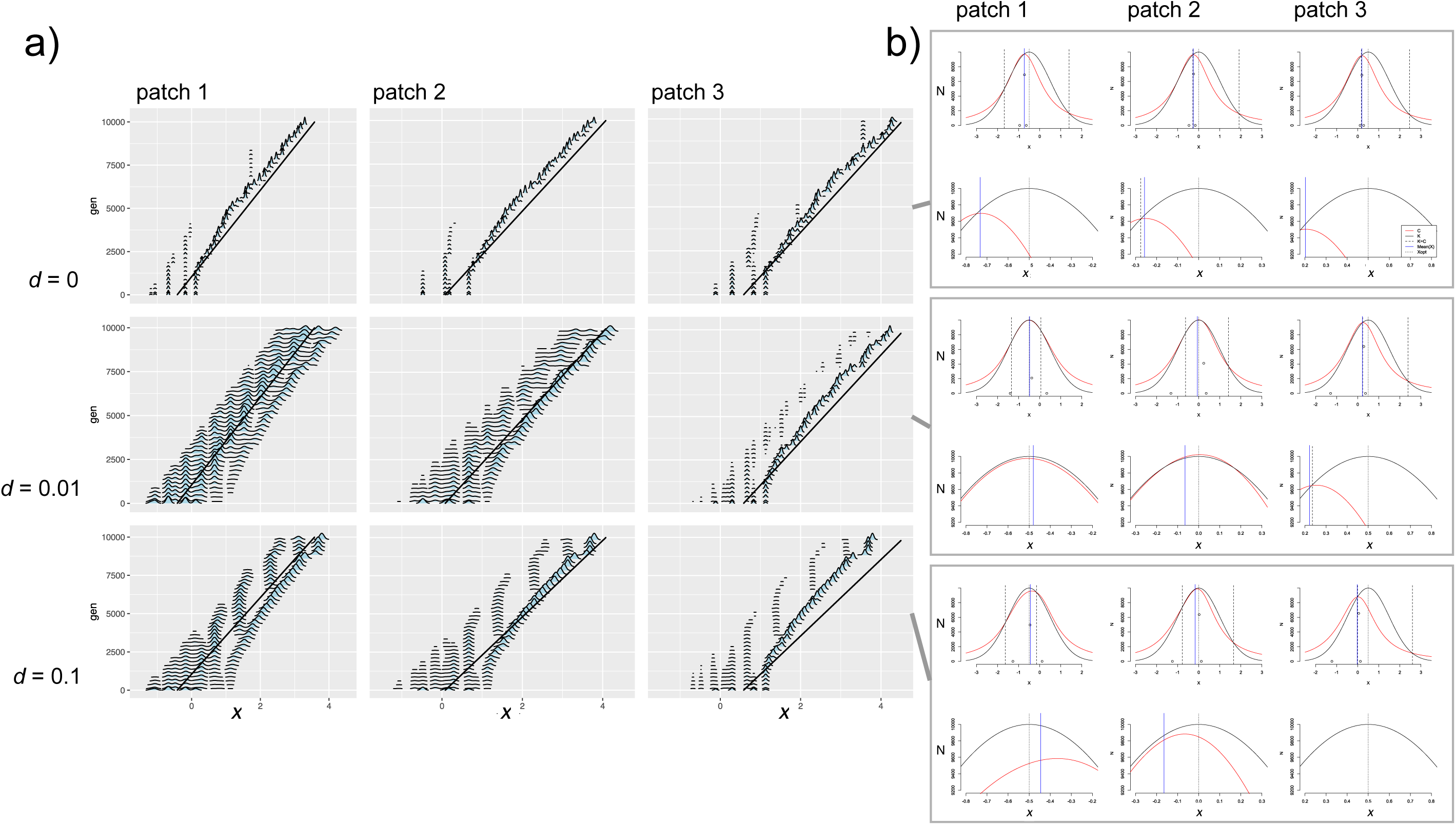
Trait evolution and competition for multi-species communities over time in a rapidly changing environment (Δ = 4×10^-4^, σ_α_ = 0.68, *d* = 0, 0.01, and 0.1). (a) Distributions of phenotypes in populations and communities over 10,000 generations of the evolving metacommunity simulation (for evolution over 50000 generations, see Figure 3, S5, and S6, third row). At all levels of dispersal, all but a single species went extinct within ∼6000 generations. When *d* > 0, competition with new mutant populations leads to periodic selection for divergent trait values but these are then lost to extinction as the environment continues to change. This intraspecific diversity is always highest in patch 1, where selection for increasing traits is driven by environmental change but immigration from the other patches creates increased selection pressure for lower competition strength and thus lower trait values. (b) Competition strength (*C*(*x*), red line) and carrying capacity (*K*(*x*), black line) as a function of trait values (*x*-axis). Without dispersal, the changing environment selects for individuals with higher trait values in all patches. When patches are linked by dispersal, immigrants are introduced and in patch 1, these increase the fitness of individuals with lower trait values that experience less competition than individuals with higher trait values. These mutant populations arise periodically then become extinct as the environment continues to change. Features of all plots are as described in Figure 2.

### Spatial structure and directional environmental change

#### 1. Shifts in direction of selection by environmental optimum and competition

*Slow environmental change*: In the full model with dispersal and slow environmental change (Δ = 10^-5^), evolutionary trajectories result from the interacting effects of a shift in the resource optimum and introduction of immigrant competitors altering the local fitness landscape. Environmental change selects for individuals with higher trait values in all patches as the *K*(*x*) curve shifts to the right, while dispersal introduces competitors that increase selection against traits in the direction of the regional mean 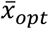 (e.g. against higher trait values in patch 1 and against lower trait values in patch 2; Figure S7). The patch-specific shift in *C*(*x*) plays a strong role in determining subsequent eco-evolutionary dynamics. In patch 1, the environment selects for individuals with higher trait values, while immigrants with high trait values from patch 2 and 3 lead to reduced competition and increased selection for individuals with lower trait values. The resulting region of trait space where *K* > *C* thus varies as a result of dispersal rate and speed of evolutionary change. With slow environmental change, individuals with lower trait values are selected for in patch 1 (Δ = 10^-5^, *d* = 0.01 and *d* = 0.1; Figures S5, S6, S7). In patch 3, directional environmental trait selection and selection for reduced competition are in the same direction, favoring evolution of increasing trait values (Figures S5, S6, S7). The mean of the phenotypic distribution in patch 2 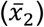 is similar to the patch optimum *x*_*2,opt*_, indicating that species in the metacommunity are able to track the slow environmental change. Although the environmental change is gradual enough that fitness landscapes are similar when comparing Δ = 0 and 10^-5^, the gradually increasing optimal trait values are still reflected in the selection for some individuals with higher trait.

#### Fast environmental change

When environmental change is more rapid (Δ = 4×10^-4^), among-patch variation in the fitness landscape increases with increasing dispersal. When *d* = 0.01, in patch 1, environmental selection for individuals with increased trait values differs from the direction of selection for individuals with lower trait values and decreased competition, leading to a wide range of species with positive carrying capacity. In patch 3, selection towards increased trait values favored by the changing environment is increased further by the reduced competition these individuals experience (Figure 4b). At the highest level of dispersal, patch 1 selects more strongly for individuals with decreased trait values, while patches 2 and 3 experience selection for increased trait values.

This among-site variation in the direction of selection creates a dynamic feedback loop between evolutionary dynamics in Patch 3 and ecological dynamics in patch 1 - the increasing environmental optimum trait value selects for individuals with increasingly higher trait values in patch 3, and these individuals immigrate into patch 1, where they increase competition for individuals with high trait values, which in turn creates selection in favor of individuals with lower trait values. These individuals then immigrate into patch 3 and combine with environmental change to increase the strength of selection for individuals with higher trait values in that patch (Figure 4b).

### 2. Phenotypic distributions, diversity, stability, and degree of maladaptation

#### Slow environmental change

Observed trajectories of evolutionary and ecological properties reflect that fitness landscapes vary in time and space due to the combination of directional environmental selection and competition strength. With slow environmental change in the absence of dispersal (Δ = 10^-5^, *d* = 0), adaptive trait shifts primarily occurred in resident species with the highest trait value and species with low trait values declined to extinction (and thus resident species traits showed a matching to the local environmental optima, indicating species sorting). This differed when *d* > 0, as the increased selection for higher trait values in patch 3 creates immigrants that inhibit adaptive directional trait evolution in resident species with high trait values in patch 1 and 2 (Figure 3 vs. S5). These immigrants also increase the strength of competition for species with high trait values, increasing selection for individuals with lower trait values in patches 1 and 2 (and thus resident species show less of a match to the local environmental optima, reducing the pattern of species sorting). In these patches, species with lower trait values can thus persist for longer periods of time and respond to environmental change by evolving increased trait values in some instances (e.g. in patch 1; Figure S5). As a result, species diversity is not lost with increasing dispersal levels as is the case when *d* = 0 (Figure S8). At the highest dispersal level, adaptive directional trait evolution is limited to the species with the highest trait value in patch 3 but a diverse species assemblage persists for the 50,000 generations considered for this (Figure S5, S6). SSD and CV both increased with increasing dispersal (Figure S10, Figure S11).

#### Fast environmental change

Dynamics under rapid environmental change (*d* > 0, Δ = 4×10^-4^) differed substantially. Species diversity is rapidly lost (Figure 4a; Figure S5; Figure S6). Only the species in the metacommunity with the highest trait value adapts, as novel mutants from other species experience increased competition and are lost to extinction. Due to dispersal, patch 3 immigrants colonize patches 1 and 2 and their competition shifts selection towards favoring individuals with lower trait values in these patches (the eco-evolutionary feedback loop described previously). However, selection against these species by the increasing environmental optimum is still stronger than for reduced competition (Figure 4b) and species with lower trait values are lost to extinction within ∼5000 generations (Figure 4a; Figure S5, Figure S6). The domination of the entire metacommunity by one species reflects a pattern of metacommunity monopolization.

The very broad region in trait space where *K* > *C* creates particular evolutionary dynamics (Figure 4b). Individuals with a broad range of trait values can persist for long periods of time, until they are outside the survivable range of *K* > *C* and eventually go extinct. As a result, though a single species from patch 3 monopolizes the entire metacommunity (Figure S8), diverse lineages persist in patch 1 and 2 and maintain an intraspecific competitive hierarchy (Figure 4a; Figure S9). This is observed most strongly at the intermediate dispersal level (middle row of Fig. 4a). With higher dispersal, competition and directional environmental change combine for a strong selection pressure favoring higher trait values in patch 3, and the monopolizing species diminishes the maintenance of genetic diversity in the other patches (though some novel lineages that are favored for transient periods of time with reduced competition develop in patch 1 and 2; Figure 4a; Figure S6; Figure S9). The among-patch variation in fitness landscapes created by the combination of environmental change and dispersal also leads to among-patch variation in lagging behind the local environmental optimum trait value. When *d* = 0.01, the mean phenotype in patch 1 matches *x*_1,*opt*_ closely and when *d* = 0.1, the population mean trait value exceeds the local optimum, indicating that immigration also reduces the lag in adaptation otherwise observed in many simulation conditions (Figure 4). SSD increased with dispersal and its variance over time increased with rapid environmental change (Figure 10). Rapid environmental change led to strong among-patch variation in CV (Figure S11).

#### Increasing niche width

Increasing niche width (σ_α_ = 0.85, 1.5) means that mutants with increasingly divergent trait values experience stronger competition than when σ_α_ = 0.68 (Figure 1b). As a result, fewer species coexist in a patch (two at relatively high abundance and one at low abundance) in the absence of dispersal when σ_α_^2^ = 0.85 and only one to two (the second at low abundance) species per patch can exist for σ_α_^2^ = 1.5 (see *d* = Δ = 0; Figure 5a; Figure S14, S15, S16, Figure S21, S22, S23, S17, S24). However, dispersal introduces individuals with very different trait values and as a result the *C*(*x*) curve has a wider area for more diverse mutants to persist (Figure 5b; Figure S12), meaning that intraspecific diversity for resident species is higher when *d* > 0 (i.e. increased width of the phenotypic distribution for resident species Figure 5a and 5b; Figure S18, Figure S25). As was observed for σ_α_ = 0.68, intraspecific diversity was highest at the intermediate dispersal level (*d* = 0.01), caused by the wide range of trait values with positive carrying capacity (Figure S12). Patches had highest local α diversity at the highest dispersal level (Figure S17, S24).

**Figure 5.**
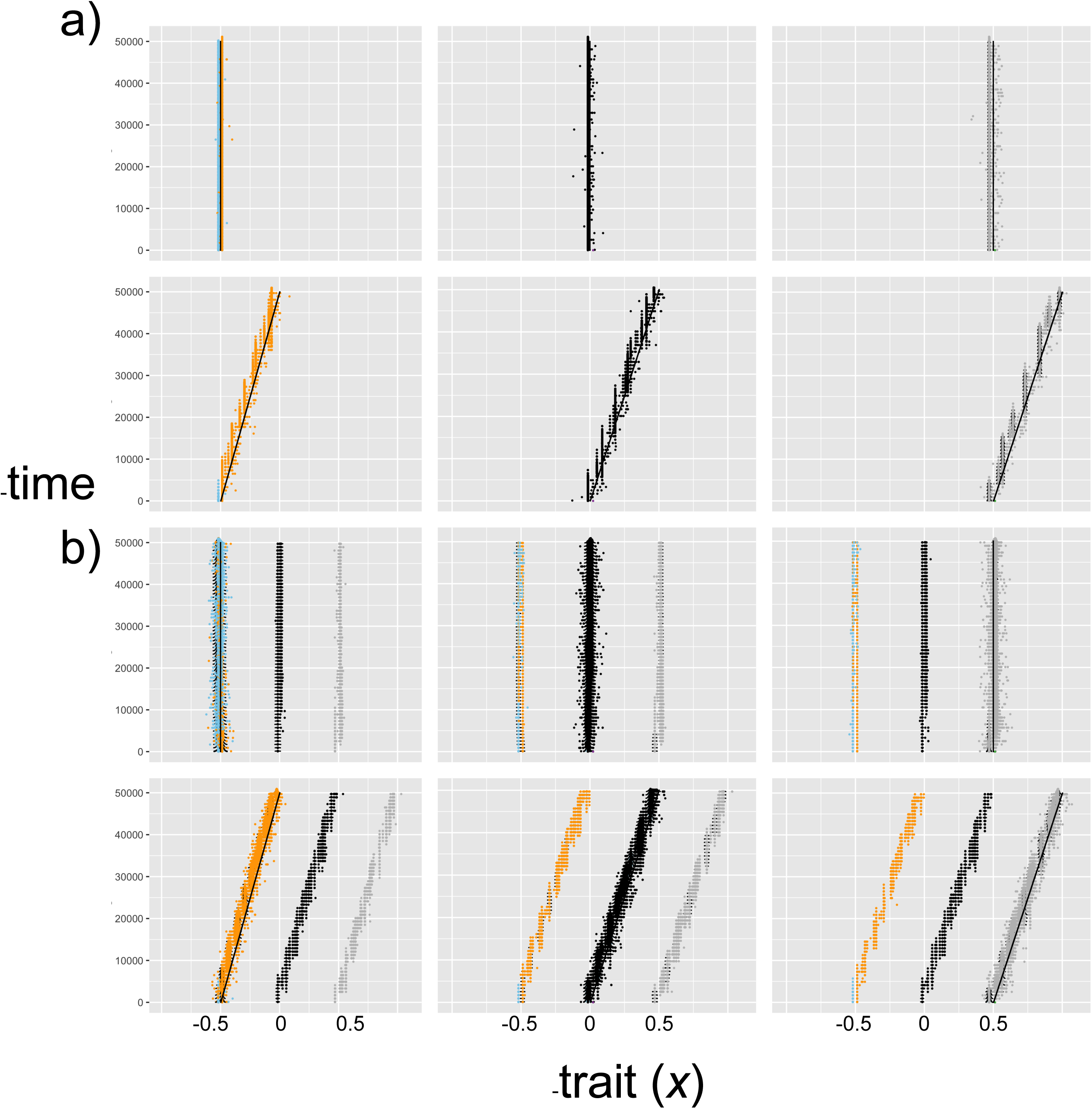
Impacts of competition strength, dispersal rate, and speed of environmental change for community trait distributions over time. Plots give phenotypic distributions (*x*-axis is trait value, height of density plots is relative abundance of individuals with that trait value, *y*-axis is time; the time series are thinned to show every 800 generations) and the solid black line in each plot indicates the local optimum trait value over time. (a) When niche width is high (σ_α_ = 1.5), in the absence of dispersal (*d* = 0) fewer species exist because individuals with similar trait values face strong competition. These species can adapt to slow environmental change (upper row: Δ = 0; lower row: Δ = 10^-5^) but retain little intraspecific diversity. (d) Dispersal (*d* = 0.1; σ_α_ = 1.5; upper row: Δ = 0; lower row: Δ = 10^-5^) not only introduces more interspecific diversity (as species from other patches have distinct enough trait values that they can persist in other patches but at lower population sizes), it also increases the amount of intraspecific diversity, as competition with other species increases the range of viable mutants that can persist with positive population size away from the local environmental optimum.

With the intermediate niche width (σ_α_ = 0.85), slow environmental change (Δ = 10^-5^) caused species with similar trait values to be lost to extinction, but after that, both remaining species were able to adaptively track environmental change (Figure S14). There was enough phenotypic space between the two remaining species that the species in each patch with the highest trait value was able to undergo branching, where a population with trait values lower than the adaptive optimum but with reduced competition intensity developed and persisted (Figure S14; Figure S13). However, when dispersal was also introduced (*d* > 0, Δ = 10^-5^), immigrants with higher trait values colonized each patch and inhibited adaptive diversification of both of the resident species (Figure S15, S16). Despite this inhibition of evolution towards the environmental optimum, however, the species with the lowest trait value in patch 1 maintained genetically diverse populations when *d* = 0.01, Δ = 10^-5^ (seen in the width of the phenotypic distribution for species colored orange in Figure S15). This resulted because the combination of selection pressures for competition from immigrants and for directional environmental change left a very wide region of trait space with positive carrying capacity values (Figure S13). At fast speed of environmental change (Δ = 4×10^-4^), species diversity was again reduced to a single species (which maintains high genetic diversity, with the highest level at intermediate dispersal level; Figure S14, S15, S16 and Figure S18; for SSD see Figure S19; for CV see Figure S20).

Dynamics differed slightly when σ_α_ = 1.5 because fewer species were able to coexist in the system, but the same general patterns were observed, where environmental change and dispersal introduced increased genetic diversity, and the species were able to track the adaptive optimal trait value. This tracking of the environmental optimum was uninhibited by interspecific competition (Figure 5; Figure S21, S22, S23; for SSD see Figure S26; for CV see Figure S27).

## Discussion

### Overview

Species that inhabit changing environments do not live in isolation. Interactions with other species can alter adaptive trajectories (Barraclough 2015), and phenotypic traits that influence fitness in the context of the abiotic environment can also be influenced by biotic interactions (Voje et al. 2015). Incorporating phenotypic evolution, species interactions, and movement in spatially variable landscapes are critical components of models needed to predict how biodiversity will respond to environmental change in the Anthropocene (Urban et al. 2016). We used a simulation model to evaluate how competition and spatial structure influence species responses to directional environmental change. We found three main results. First, spatial structure introduced among-site variation in the direction of selection. In some sites trait evolution towards the adaptive optimum (i.e. the resource maximum) was accelerated as selection from the directionally changing environment and from competition worked in the same direction, while at other sites competition inhibited local adaptation. Second, metacommunity patterns such as species sorting (the match between species traits and the local environment) and monopolization (where diversity is lost and one genotype or species dominates the landscape) differ from those observed in previous models that do not treat phenotype fitness as a function of both the local environment and competition with other individuals. Species sorting can be diminished even at a low dispersal rate, as selection favors species with lower carrying capacity but reduced competition, meaning that not all species will evolve towards the environmental optimum trait value. Monopolization occurs at a fast rate of environmental change, but intraspecific phenotypic diversity was maintained even in this situation and thus genotypic monopolization was not observed. Third, we observed an eco-evolutionary feedback loop between local phenotypic evolution at one site and competition at another site. This feedback loop was maintained by dispersal and indicates that eco-evolutionary dynamics in communities operate at a landscape scale.

### Dynamic variation in fitness landscapes over time and space

Our results at the local scale reflect the findings of two previous studies that modeled trait evolution and competition using an adaptive dynamics framework, and builds on them by considering evolution and competition in a spatially structured landscape. Johansson (2008) showed that multiple competing species are retained under slow environmental change and that species are increasingly lost to extinction under fast environmental change. Our model, which begins from the same model Johansson (2008) used, thus repeats these results (Figure 3). We also show that shifts in the fitness landscape caused by rapid environmental change select only for individuals with increasingly higher trait values (and therefore species with lower trait values are lost even though they experience decreased competition; Figure 4, *d* = 0). By introducing movement among patches, we found that persistent immigration of competitors also shifts the fitness landscape, in a way that increases selection for individuals with low trait values in patch 1 and high trait values in patch 3 (Figure 4). The evolutionary dynamics observed in the presence of both dispersal and environmental change are therefore a product of variation in the fitness landscape over time and space. Osmond & de Mazancourt (2013) showed how variation in the shapes and positions of the carrying capacity function *K*(*x*) and the competition function *C*(*x*) determines the speed and direction of phenotypic evolution. Our results simply introduce spatial structure as one mechanism by which the direction and magnitude of selection vary among populations. The competition curve is not fixed, but is instead frequency dependent, and movement of individuals among patches shifts the frequency distribution of trait values. The *C*(*x*) curve thus dynamically shifts as dispersal introduces immigrants that alter local phenotypic and population size distributions and, as a consequence, the amount of competition experienced by individuals. Our study presents a first step towards characterizing how landscape features and connectivity patterns can drive eco-evolutionary dynamics in multi-species communities.

### Metacommunity patterns for trait and species diversity

Previous theoretical studies have considered how environmental variation influences trait evolution in communities of competitors in a spatially explicit landscape. In these studies, phenotypic traits were assumed to evolve in response to environmental variation (among sites: de Mazancourt et al. 2008; Loeuille & Leibold 2008; Urban & De Meester 2009; Vanoverbeke et al. 2016; across sites and over time: Norberg et al. 2012). Species assemblages emerged as a result of environmental heterogeneity and site connectivity. Maladaptation (a mismatch between species traits and the local environment) could result due to spatial mass effects, low levels of genetic variance reducing adaptive evolution, or combinations of dispersal and amounts of genetic variation (i.e. the ‘global monopolization’ of one or a few species due to rapid adaptation or evolutionary priority effects). Our model differs from these studies by considering that competition depends on the phenotypic distance between interacting individuals (using competition coefficients α_*ij*_ instead of competition for microsites within patches) and by modeling evolution in a phenotypic trait that influences both local carrying capacity and strength of competition (instead of in a trait that influences only a component of population dynamics such as recruitment, survival, or growth). Considering evolution of traits that reflect fitness tradeoffs and facilitate coexistence (e.g. Vasseur et al. 2011; Kremer & Klausmeier 2013) is an important step to better understand the dynamics of trait and species composition in nature.

Modeling competition as a function of the trait that also determines carrying capacity revealed some differences compared to previous studies of evolving metacommunities. In the slowly changing environment (Δ = 10^-5^), when sites were not connected by dispersal, at least one species in each patch showed adaptation to the local optimal carrying capacity (Figure 3b). However, when dispersal was included, fewer species could adapt to the local environment due to competitive constraints, and thus species sorting (i.e. the degree of matching between resident species traits and the local environmental optimum) was diminished over time (Figure S5; also reflected in the increasing sum of squared deviations from the optimal trait value with dispersal seen in Figures S1 and S3). This matches the results of Norberg et al. (2012), who also modeled competition directly using interaction coefficients. They found that species extinctions were hastened with increasing dispersal and species sorting was diminished, whereas the study of Loeuille & Leibold (2008) found that species sorting was maintained in a lottery competition model when dispersal was high relative to evolutionary rates. Our study took an additional step of varying the strength of competition (i.e. the phenotypic range over which individuals experience competition; Figure 1b). We found that the inhibition of species sorting depended on niche width, where metacommunities with narrower niche widths were more likely to maintain patterns of species sorting even under high dispersal (Figure 5; also reflected in the greatly reduced deviations from optimal trait values observed in Figure S26). Like the previous studies of evolution in metacommunities, we observed global monopolization where one species dominates the landscape (Loeuille & Leibold 2008; Urban & De Meester 2009; Vanoverbeke et al. 2016), but in our system this depended on the speed of environmental change, and thus on the magnitude of the directional environmental selection pressure. Loeuille & Leibold (2008) observed global monopolization of the metacommunity by a single species at a combination of high dispersal and high mutation rates. While we did not vary the mutation rate, it is possible that the rate of environmental change played a similar role. Our results also indicate that the monopolizing species can maintain very high levels of intraspecific phenotypic diversity especially at our intermediate dispersal rate (Figure 3, 4; Figures S5, S6). In the future, models of evolving metacommunities can benefit from considering additional factors such as the evolution of multiple correlated traits or varying genetic variance (Edwards et al. 2018) and response to multiple environmental stressors (Jackson et al. 2015).

### Eco-evolutionary feedback loops and adaptive dynamics

Trait evolution in species that compete for resources and inhabit a metacommunity is influenced by intriguing mechanisms that were previously detailed in the fields of adaptive dynamics and eco-evolutionary dynamics. Eco-evolutionary feedback loops are cyclical interactions between ecology and evolution, where changes in ecological interactions drive evolutionary change in traits that, in turn, feed back to alter the ecological interactions in a continuing process (Post & Palkovacs 2009). These feedback loops have been demonstrated empirically for population, host-parasite and predator-prey, and ecosystem dynamics in closed systems (e.g. Yoshida et al. 2003; Turcotte et al. 2013; Matthews et al. 2016; Brunner et al. 2017) and for evolution of dispersal and metapopulation dynamics in spatially structured populations (Fronhofer & Altermatt 2015; Legrand et al. 2017). There is some evidence that eco-evolutionary feedbacks can facilitate coexistence (terHorst et al. 2018), but these are not well-studied at the metacommunity scale (but see Colombo et al. 2018 for a model of predator-prey interactions in a spatially structured system). We identified an eco-evolutionary feedback loop in our system - evolution in patch 3 selects for individuals with higher trait values, immigrants from patch 3 with high trait values shift competition strength in patch 1 and select for individuals with lower trait values, and then immigrants from patch 1 with lower trait values move to patch 3 where they complete the feedback loop, increasing selection for higher trait values. This effect was observed at all combinations of dispersal and environmental change rate > 0 but is easiest to visualize at the highest levels of both (Figure 4b). This is a novel mechanism for an eco-evolutionary feedback that involves dispersal maintaining dynamic reinforcement between competition and phenotypic evolution.

Adaptive dynamics theory is well suited to model eco-evolutionary dynamics because the equations explicitly link trait evolution to ecological processes and have been used to understand the process of evolutionary rescue (where initially maladapted populations adapt to the environment and avoid extinction, Gomulkiewicz & Holt 1995; e.g. Ferriere & Legendre 2013). Our simulation model reproduced expected elements observed in previous adaptive dynamics models, such as the branching process that produced evolutionarily stable trait values for species generated in our initialization step. The same branching process produced the divergence of populations into multiple coexisting sub-populations within a site observed in some conditions, such as in patch 3 when the rate of environmental change is slow and sites are linked by dispersal (final 10,000 generations in Figure 3, Figure S5). Adaptive dynamics models have also revealed processes where populations can evolve towards either their own extinction (evolutionary suicide, Ferriere 2000; Gyllenberg & Parvinen 2001; Gyllenberg et al. 2002) or between two equilibria, one at low and one at high population density (evolutionary collapse, Dercole et al. 2002). In both instances, selection can produce mutants with traits that inhabit a trait space beyond the extinction boundary, where the mutant population cannot persist even though it can be maintained for long periods of time. In some conditions, our model reflected the long-term persistence but eventual extinction of mutants in the population. For example with σ_α_ = 0.68, *d* = 0.1, and Δ = 4×10^-4^ (Figure 4), a primary resident population in patch 3 tracked the local environmental optimum, but immigrants from patch 3 shifted the local competition curve in other patches in a way that permitted mutant individuals with a very wide range of trait values to persist for long periods of time before declining to extinction. Figure 4 shows that populations branch from the main resident population and survive, decline to extinction, and the process repeats itself. Although this process is likely not the same as the mechanism that produces suicide and collapse, it indicates that mal-adaptive diversity can be maintained for relatively long periods of time.

### Conclusions

As species encounter changing climates, it is critical to understand whether they will adapt, move to more suitable habitats, or instead decline to extinction. Models of species range shifts are increasingly considering genetic variation, evolution, and species interactions (e.g. Bocedi et al. 2014), and species distribution models increasingly include species association matrices as well as local adaptation and phenotypic plasticity (Wisz et al. 2013; Pollock et al. 2014; Benito Garzón et al. 2019). Accurate representations of these more complex processes will be necessary to forecast how biodiversity will track climate change (Urban et al. 2016). Our study considers how trait evolution in metacommunities can respond to environmental change and it adds to previous studies by considering phenotypic evolution in a trait that represents a balance between two selection pressures - the changing environment and the strength of competition with other populations and species consuming a similar range of resources. The placement of the evolving community in a landscape introduced feedbacks between eco-evolutionary processes that are driven by dispersal and that maintain biodiversity despite the increased connectivity of species among sites. One possible consequence of habitat fragmentation or loss may be the disruption of these spatial eco-evolutionary feedbacks and a resulting loss of biodiversity.

## Supplementary Materials

The following are available online at www.mdpi.com/xxx/s1, Supplementary Information, Figure S1-S27.

## Author Contributions

Conceptualization, J.P.; methodology, J.P. and A.F.; software, J.P. and A.F.; validation, J.P. and A.F.; formal analysis, J.P. and A.F.; investigation, J.P. and A.F.; resources, J.P. and A.F.; data curation, J.P. and A.F.; writing—original draft preparation, J.P.; writing—review and editing, J.P. and A.F.; visualization, J.P. and A.F.; supervision, J.P.; project administration, J.P.; funding acquisition, J.P. and A.F. All authors have read and agreed to the published version of the manuscript.

## Acknowledgements

We would like to thank N. Loeuille for his guidance in formulating research questions. The authors acknowledge William & Mary Research Computing for providing computational resources and/or technical support that have contributed to the results reported within this paper. URL: https://www.wm.edu/it/rc. APF was supported by the William & Mary 1693 Scholars Program. JHP acknowledges support from the The Richard Lounsbery Foundation, for the project *Urbanization and land use change effects on aquatic biodiversity*.

## Supplementary Code

Code will be made available on the GitHub page of J.H. Pantel at time of publication.

## Supplementary Information for Eco-evolutionary feedbacks and the maintenance of metacommunity diversity in a changing environment

**Figure S1**. Inter- and intraspecific diversity, deviation from optimum trait value, and coefficient of variation over time in an unchanging environment (Δ = 0, σ_α_ = 0.68, *d* = 0, 0.01, and 0.1). Row 1 is interspecific diversity (grey: α, black: γ, orange: β). Row 2 is intraspecific diversity, where every color is a species (corresponding to the colors in Figure 2). Row 3 is the sum of squared deviations of trait values from the local environmental optimum trait value for all individuals in a patch (black: patch 1, red: patch 2, blue: patch 3). Row 4 is the coefficient of variation of total population size across all species at the local (patch 1: grey, patch 2: light blue, patch 3: orange) and metacommunity (black) level.

**Figure S2**. Competition, carrying capacity, and population trait values in a changing environment. In both plots σ_α_ = 0.68 and *d* = 0, in (a) Δ = 10^-5^, and in (b) Δ = 4×10^-4^. Competition functions (*C*(*x*), red), carrying capacity functions (*K*(*x*), black), population (black points), and all other features are as given in Figure 2.

**Figure S3**. Inter- and intraspecific diversity, deviation from optimum trait value, and coefficient of variation over time in a slowly changing environment (Δ = 10^-5^, σ_α_ = 0.68) across a range of dispersal levels (from left to right, *d* = 0, 0.01, and 0.1). Features of all plots are as described in Figure S1.

**Figure S4**. Phenotypic distributions over time when competition strength is low (σ_α_ = 0.68) and there is no dispersal (*d* = 0), across a range of rates of environmental change (Δ = 0, 10^-5^, and 4×10-4 from top to bottom), for 15,000 generations (the time series are thinned to show every 250 generations). This figure contains the same results as Figure 3, but the limits of the *x*-axes are the same across all the levels of environmental change, to better visualize the intraspecific diversity across conditions. Columns represent three patches across dispersal levels. Features of plots are as described in Figure 2.

**Figure S5**. Phenotypic distributions over time when competition strength is low (σ_α_ = 0.68) and there is intermediate dispersal (*d* = 0.01), across a range of rates of environmental change (Δ = 0, 10^-5^, and 4×10^-4^ from top to bottom), for 50,000 generations (the time series are thinned to show every 800 generations). Columns represent three patches across dispersal levels. Features of plots are as described in Figure 2. The nine plots with color (columns 4, 5, and 6) have the same information as the nine plots without color (columns 1, 2, and 3), and both are shown so that both intra- and interspecific diversity are visible.

**Figure S6**. Phenotypic distributions over time when competition strength is low (σ_α_ = 0.68) and there is high dispersal (*d* = 0.1), across a range of rates of environmental change (Δ = 0, 10^-5^, and 4×10^-4^ from top to bottom), for 50,000 generations (the time series are thinned to show every 800 generations). Columns represent three patches across dispersal levels. Features of plots are as described in Figure 2. The nine plots with color (columns 4, 5, and 6) have the same information as the nine plots without color (columns 1, 2, and 3), and both are shown so that both intra- and interspecific diversity are visible.

**Figure S7**. Competition, carrying capacity, and population trait values in a slowly changing environment. In all plots σ_α_ = 0.68 and Δ = 10^-5^. Competition functions (*C*(*x*), red), carrying capacity functions (*K*(*x*), black), population (black points), and all other features are as given in Figure 2 and vary across levels of dispersal (*d* = 0, 0.01, 0.1).

**Figure S8**. Interspecific diversity over time when competition strength is low (σ_α_ = 0.68) across a range of dispersal levels and rates of environmental change (grey: α diversity, black: γ diversity, orange: β diversity). Rows are increasing levels of environmental change (Δ = 0, 10^-5^, and 4×10^- 4^ from top to bottom) and columns are increasing levels of dispersal (*d* = 0, 0.01, and 0.1 from left to right).

**Figure S9**. Intraspecific diversity over time when competition strength is low (σ_α_ = 0.68) across a range of dispersal levels and rates of environmental change. Each color represents genetic diversity of a different species. Rows are increasing levels of environmental change (Δ = 0, 10^-5^, and 4×10^-4^ from top to bottom) and columns are increasing levels of dispersal (*d* = 0, 0.01, and

0.1 from left to right).

**Figure S10**. Sum of squared deviations of trait values from local environmental optimum trait value for all individuals in patches over time when competition strength is low (σ_α_ = 0.68), across a range of dispersal levels and rates of environmental change (black: patch 1, red: patch 2, blue: patch 3). Rows are increasing levels of environmental change (Δ = 0, 10^-5^, and 4×10^-4^ from top to bottom) and columns are increasing levels of dispersal (*d* = 0, 0.01, and 0.1 from left to right).

**Figure S11**. Coefficient of variation of total population size across all species at the local (patch 1: grey, patch 2: light blue, patch 3: orange) and metacommunity (black) level over time when competition strength is low (σ_α_ = 0.68), across a range of dispersal levels and rates of environmental change. Rows are increasing levels of environmental change (Δ = 0, 10^-5^, and 4×10^-4^ from top to bottom) and columns are increasing levels of dispersal (*d* = 0, 0.01, and 0.1 from left to right).

**Figure S12**. Competition, carrying capacity, and population trait values in an unchanging environment. In all plots σ_α_ = 0.85 and Δ = 0. Competition functions (*C*(x), red), carrying capacity functions (*K*(x), black), population (black points), and all other features are as given in Figure 2 and vary across levels of dispersal (d = 0, 0.01, 0.1).

**Figure S13**. Competition, carrying capacity, and population trait values in a slowly changing environment. In all plots σ_α_ = 0.85 and Δ = 10^-5^. Competition functions (*C*(x), red), carrying capacity functions (*K*(x), black), population (black points), and all other features are as given in Figure 2 and vary across levels of dispersal (d = 0, 0.01, 0.1).

**Figure S14**. Phenotypic distributions over time when competition strength is intermediate (σ_α_ = 0.85) and there is no dispersal (*d* = 0), across a range of rates of environmental change (Δ = 0, 10^-5^, and 4×10^-4^ from top to bottom), for 50,000 generations (the time series are thinned to show every 800 generations). Columns represent three patches across dispersal levels. Features of plots are as described in Figure 2. The nine plots with color (columns 4, 5, and 6) have the same information as the nine plots without color (columns 1, 2, and 3), and both are shown so that both intra- and interspecific diversity are visible.

**Figure S15**. Phenotypic distributions over time when competition strength is intermediate (σ_α_ = 0.85) and there is intermediate dispersal (*d* = 0.01), across a range of rates of environmental change (Δ = 0, 10^-5^, and 4×10^-4^ from top to bottom), for 50,000 generations (the time series are thinned to show every 800 generations). Columns represent three patches across dispersal levels. Features of plots are as described in Figure 2. The nine plots with color (columns 4, 5, and 6) have the same information as the nine plots without color (columns 1, 2, and 3), and both are shown so that both intra- and interspecific diversity are visible.

**Figure S16**. Phenotypic distributions over time when competition strength is intermediate (σ_α_ = 0.85) and there is high dispersal (*d* = 0.1), across a range of rates of environmental change (Δ = 0, 10^-5^, and 4×10^-4^ from top to bottom), for 50,000 generations (the time series are thinned to show every 800 generations). Columns represent three patches across dispersal levels. Features of plots are as described in Figure 2. The nine plots with color (columns 4, 5, and 6) have the same information as the nine plots without color (columns 1, 2, and 3), and both are shown so that both intra- and interspecific diversity are visible.

**Figure S17**. Interspecific diversity over time when competition strength is intermediate (σ_α_ = 0.85) across a range of dispersal levels and rates of environmental change (grey: α diversity, black: γ diversity, orange: β diversity). Rows are increasing levels of environmental change (Δ = 0, 10^-5^, and 4×10^-4^ from top to bottom) and columns are increasing levels of dispersal (*d* = 0, 0.01, and 0.1 from left to right).

**Figure S18**. Intraspecific diversity over time when competition strength is intermediate (σ_α_ = 0.85) across a range of dispersal levels and rates of environmental change. Each color represents genetic diversity of a different species. Rows are increasing levels of environmental change (Δ = 0, 10^-5^, and 4×10^-4^ from top to bottom) and columns are increasing levels of dispersal (*d* = 0, 0.01, and 0.1 from left to right).

**Figure S19**. Sum of squared deviations of trait values from local environmental optimum trait value for all individuals in a patch over time when competition strength is intermediate (σ_α_ = 0.85), across a range of dispersal levels and rates of environmental change (black: patch 1, red: patch 2, blue: patch 3). Rows are increasing levels of environmental change (Δ = 0, 10^-5^, and 4×10^-4^ from top to bottom) and columns are increasing levels of dispersal (*d* = 0, 0.01, and 0.1 from left to right).

**Figure S20**. Coefficient of variation of total population size across all species at the local (patch 1: grey, patch 2: light blue, patch 3: orange) and metacommunity (black) level over time when competition strength is intermediate (σ_α_ = 0.85), across a range of dispersal levels and rates of environmental change. Rows are increasing levels of environmental change (Δ = 0, 10^-5^, and 4×10^-4^ from top to bottom) and columns are increasing levels of dispersal (*d* = 0, 0.01, and 0.1 from left to right).

**Figure S21**. Phenotypic distributions over time when competition strength is high (σ_α_ = 1.5) and there is no dispersal (*d* = 0), across a range of rates of environmental change (Δ = 0, 10^-5^, and 4×10^-4^ from top to bottom), for 50,000 generations (the time series are thinned to show every 800 generations). Columns represent three patches across dispersal levels. Features of plots are as described in Figure 2. The nine plots with color (columns 4, 5, and 6) have the same information as the nine plots without color (columns 1, 2, and 3), and both are shown so that both intra- and interspecific diversity are visible.

**Figure S22**. Phenotypic distributions over time when competition strength is high (σ_α_ = 1.5) and there is intermediate dispersal (*d* = 0.01), across a range of rates of environmental change (Δ = 0, 10^-5^, and 4×10^-4^ from top to bottom), for 50,000 generations (the time series are thinned to show every 800 generations). Columns represent three patches across dispersal levels. Features of plots are as described in Figure 2. The nine plots with color (columns 4, 5, and 6) have the same information as the nine plots without color (columns 1, 2, and 3), and both are shown so that both intra- and interspecific diversity are visible.

**Figure S23**. Phenotypic distributions over time when competition strength is high (σ_α_ = 1.5) and there is high dispersal (*d* = 0.1), across a range of rates of environmental change (Δ = 0, 10^-5^, and 4×10^-4^ from top to bottom), for 50,000 generations (the time series are thinned to show every 800 generations). Columns represent three patches across dispersal levels. Features of plots are as described in Figure 2. The nine plots with color (columns 4, 5, and 6) have the same information as the nine plots without color (columns 1, 2, and 3), and both are shown so that both intra- and interspecific diversity are visible.

**Figure S24**. Interspecific diversity over time when competition strength is high (σ_α_ = 1.5) across a range of dispersal levels and rates of environmental change (grey: α diversity, black: γ diversity, orange: β diversity). Rows are increasing levels of environmental change (Δ = 0, 10^-5^, and 4×10^-4^ from top to bottom) and columns are increasing levels of dispersal (*d* = 0, 0.01, and 0.1 from left to right).

**Figure S25**. Intraspecific diversity over time when competition strength is high (σ_α_ = 1.5) across a range of dispersal levels and rates of environmental change. Each color represents genetic diversity of a different species. Rows are increasing levels of environmental change (Δ = 0, 10^-5^, and 4×10^-4^ from top to bottom) and columns are increasing levels of dispersal (*d* = 0, 0.01, and 0.1 from left to right).

**Figure S26**. Sum of squared deviations of trait values from local environmental optimum trait value for all individuals in a patch over time when competition strength is high (σ_α_ = 1.5), across a range of dispersal levels and rates of environmental change (black: patch 1, red: patch 2, blue: patch 3). Rows are increasing levels of environmental change (Δ = 0, 10^-5^, and 4×10^-4^ from top to bottom) and columns are increasing levels of dispersal (*d* = 0, 0.01, and 0.1 from left to right).

**Figure S27**. Coefficient of variation of total population size across all species at the local (patch 1: grey, patch 2: light blue, patch 3: orange) and metacommunity (black) level over time when competition strength is high (σ_α_ = 1.5), across a range of dispersal levels and rates of environmental change. Rows are increasing levels of environmental change (Δ = 0, 10^-5^, and 4×10^-4^ from top to bottom) and columns are increasing levels of dispersal (*d* = 0, 0.01, and 0.1 from left to right).

