## Supplementary figures and images for "Eco-evolutionary feedbacks and the maintenance of metacommunity diversity in a changing environment"

### S1

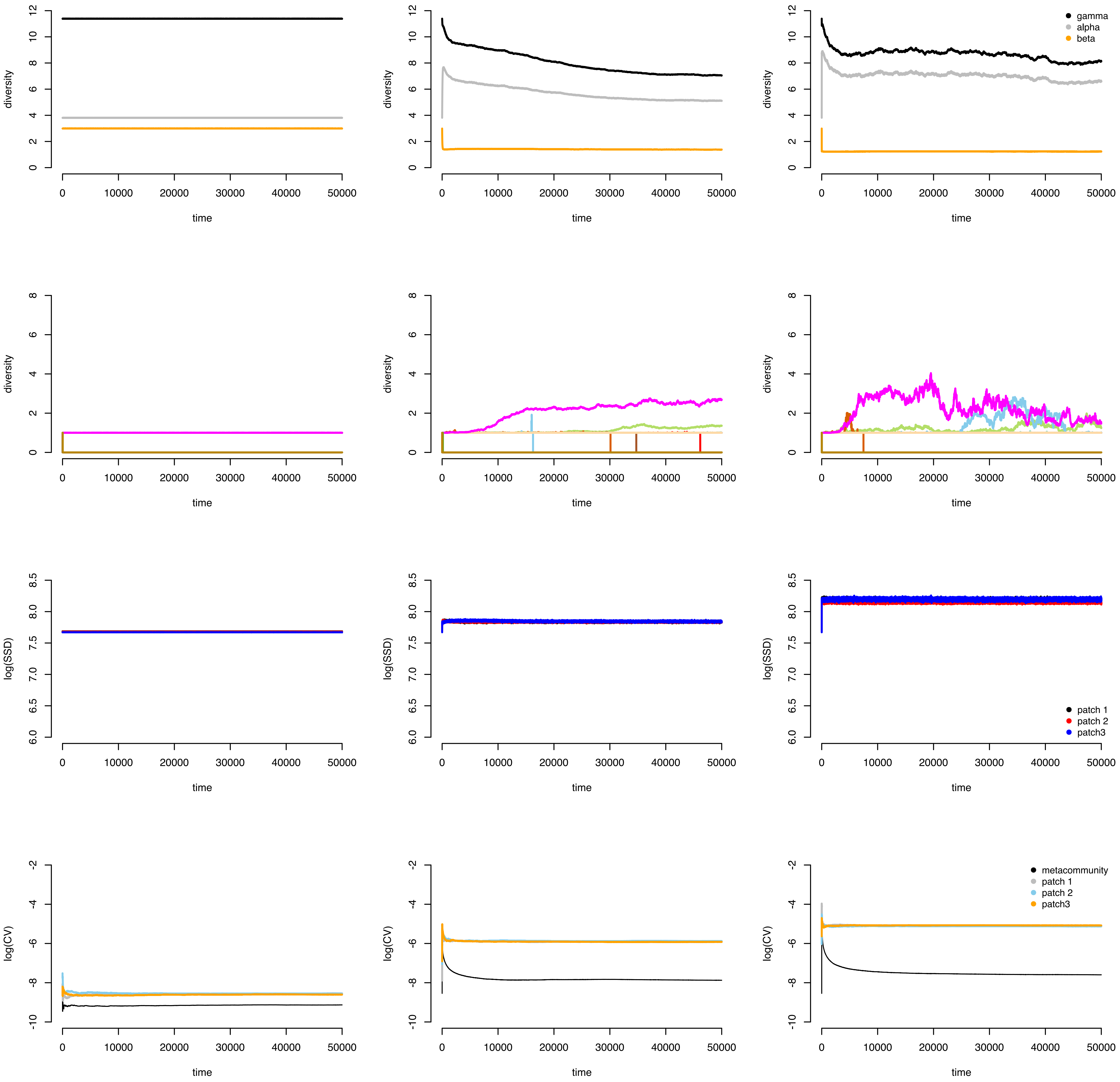

### S2

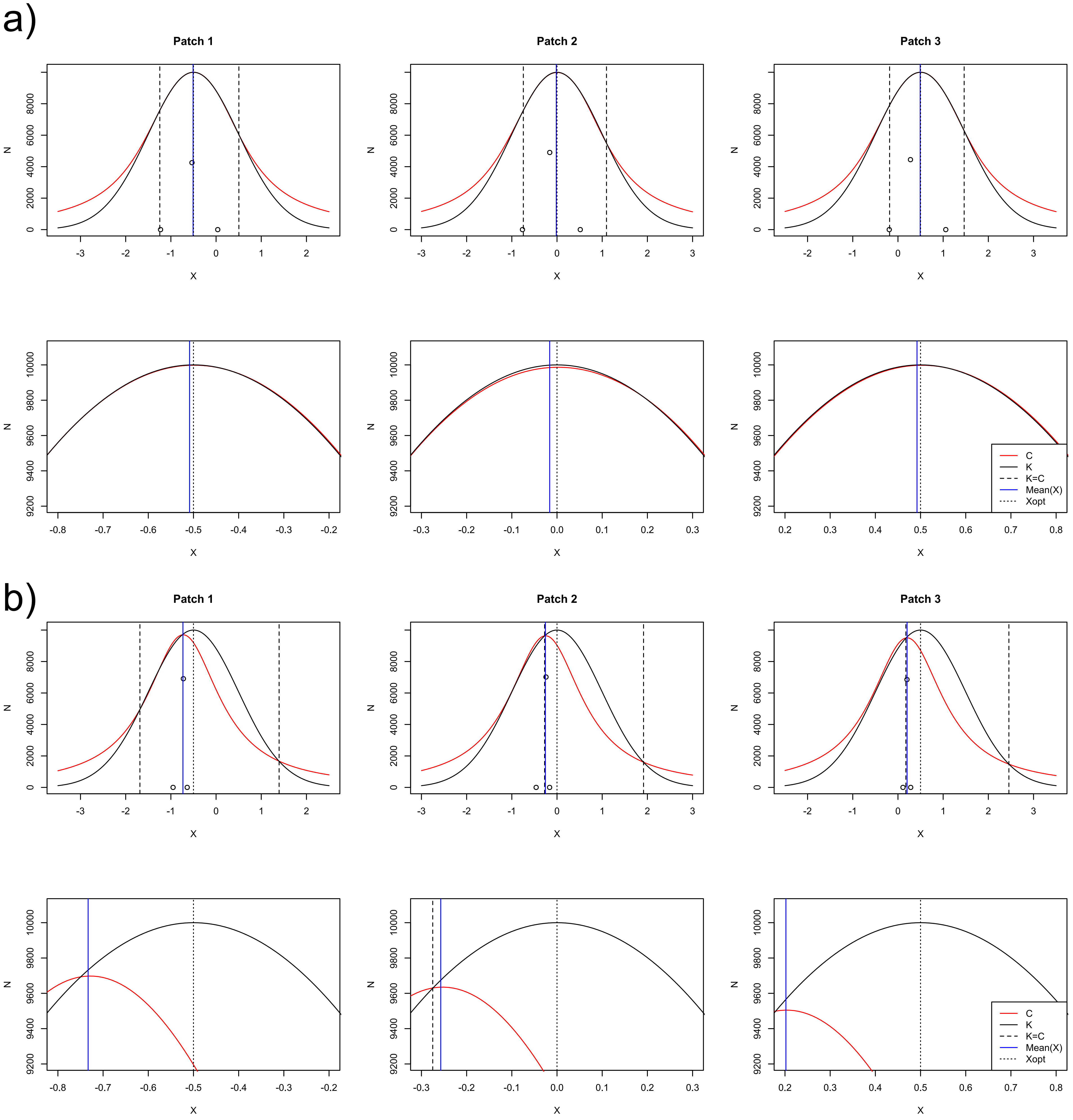

### S3

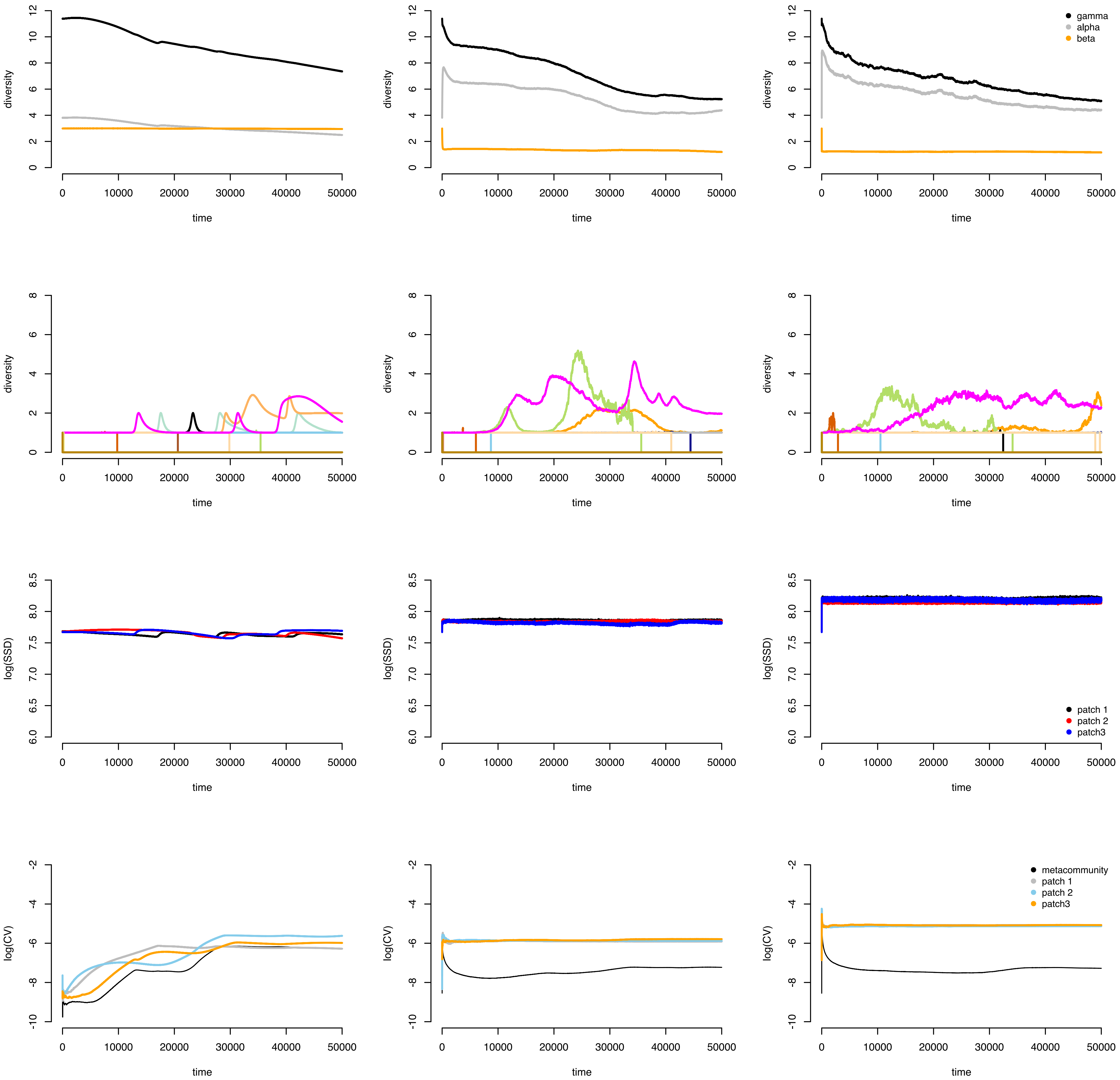

### S4

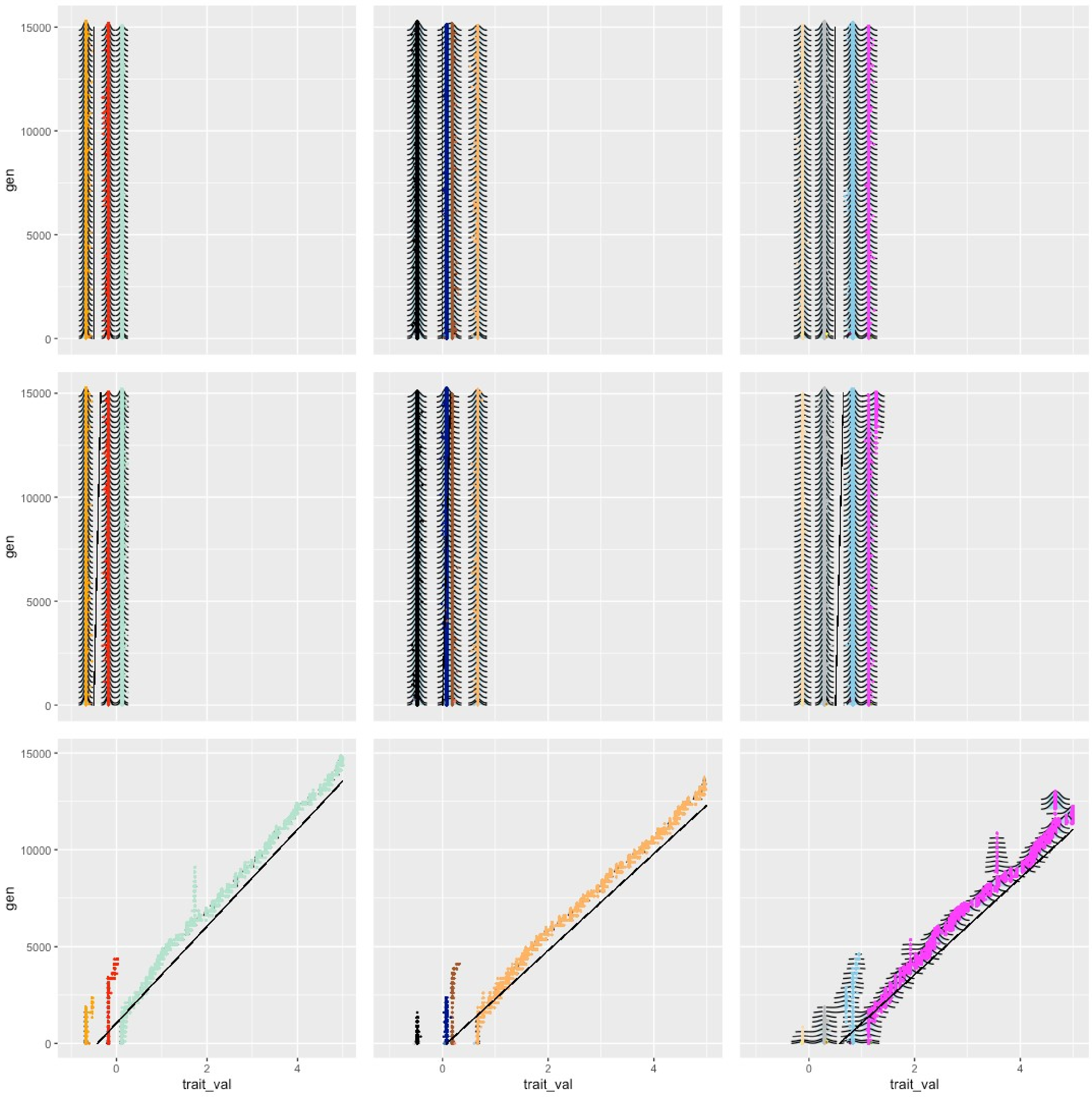

### S5

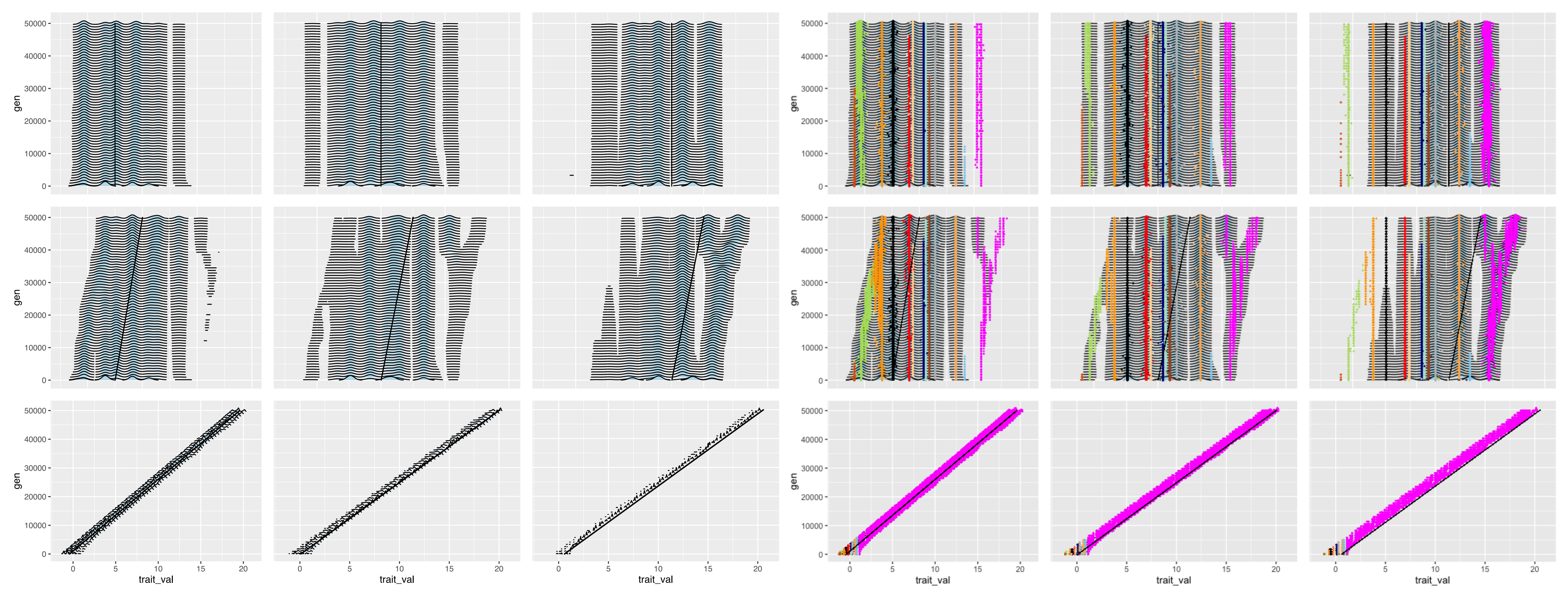

### S6

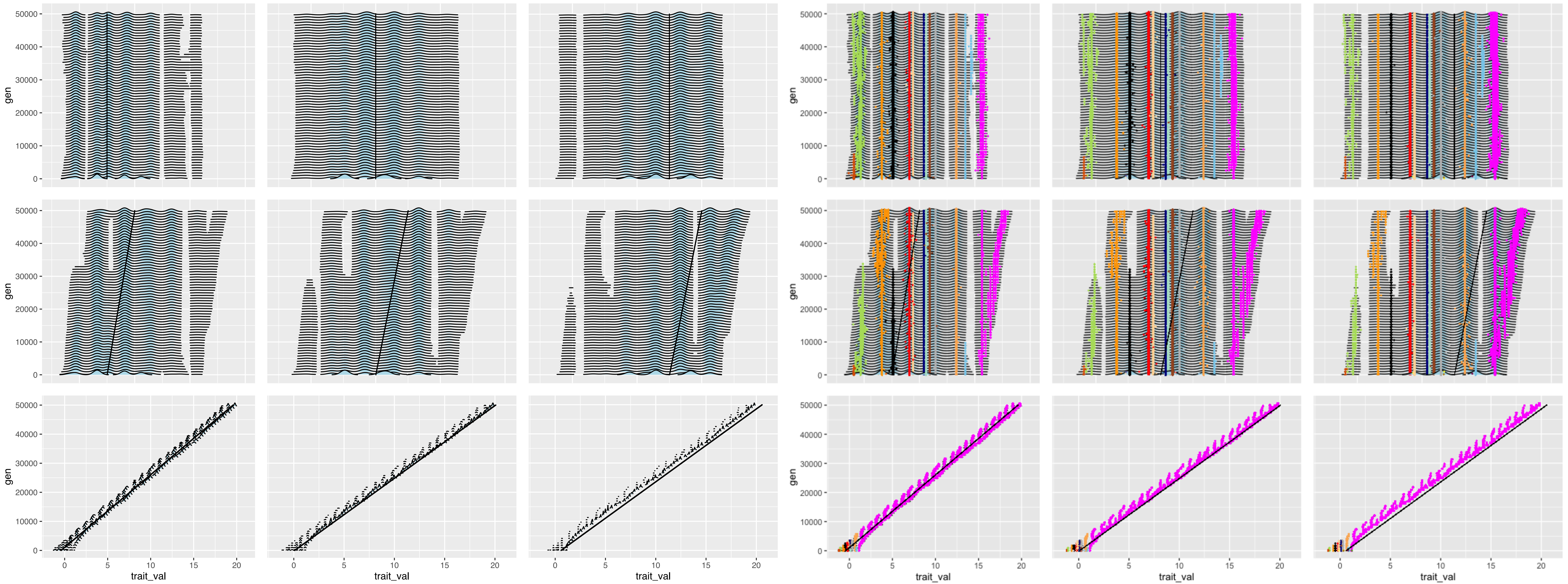

### S7

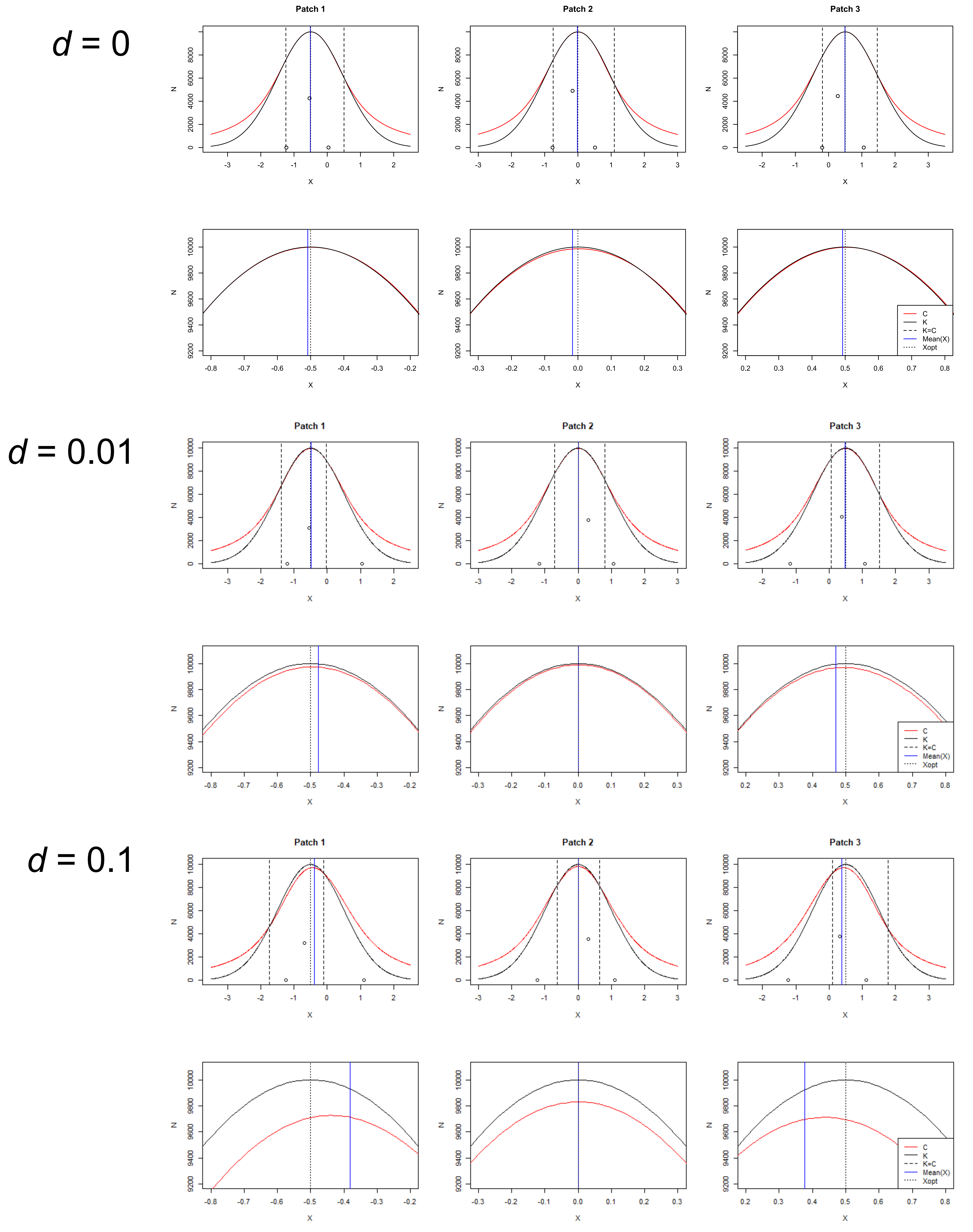

### S8

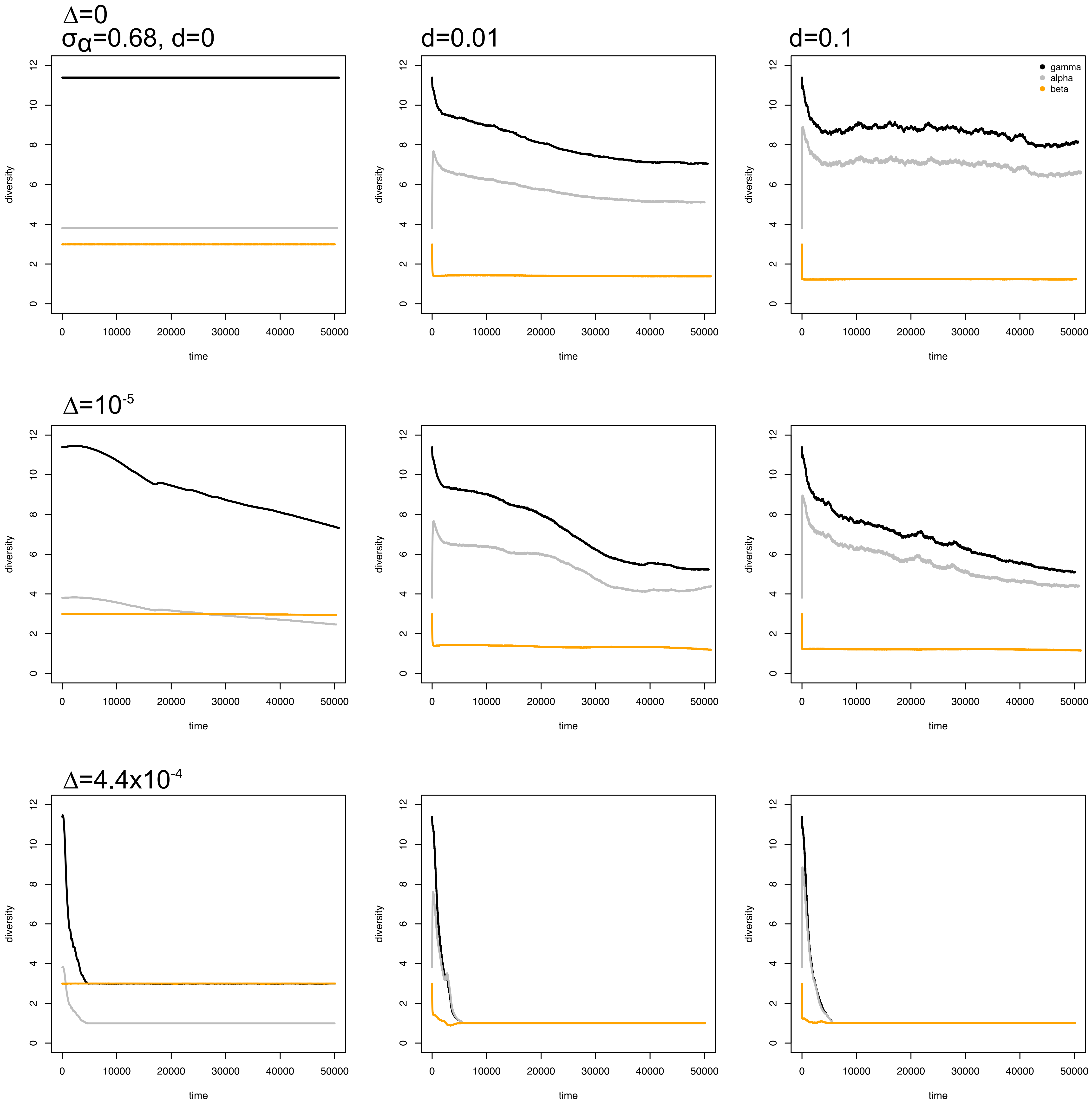

### S9

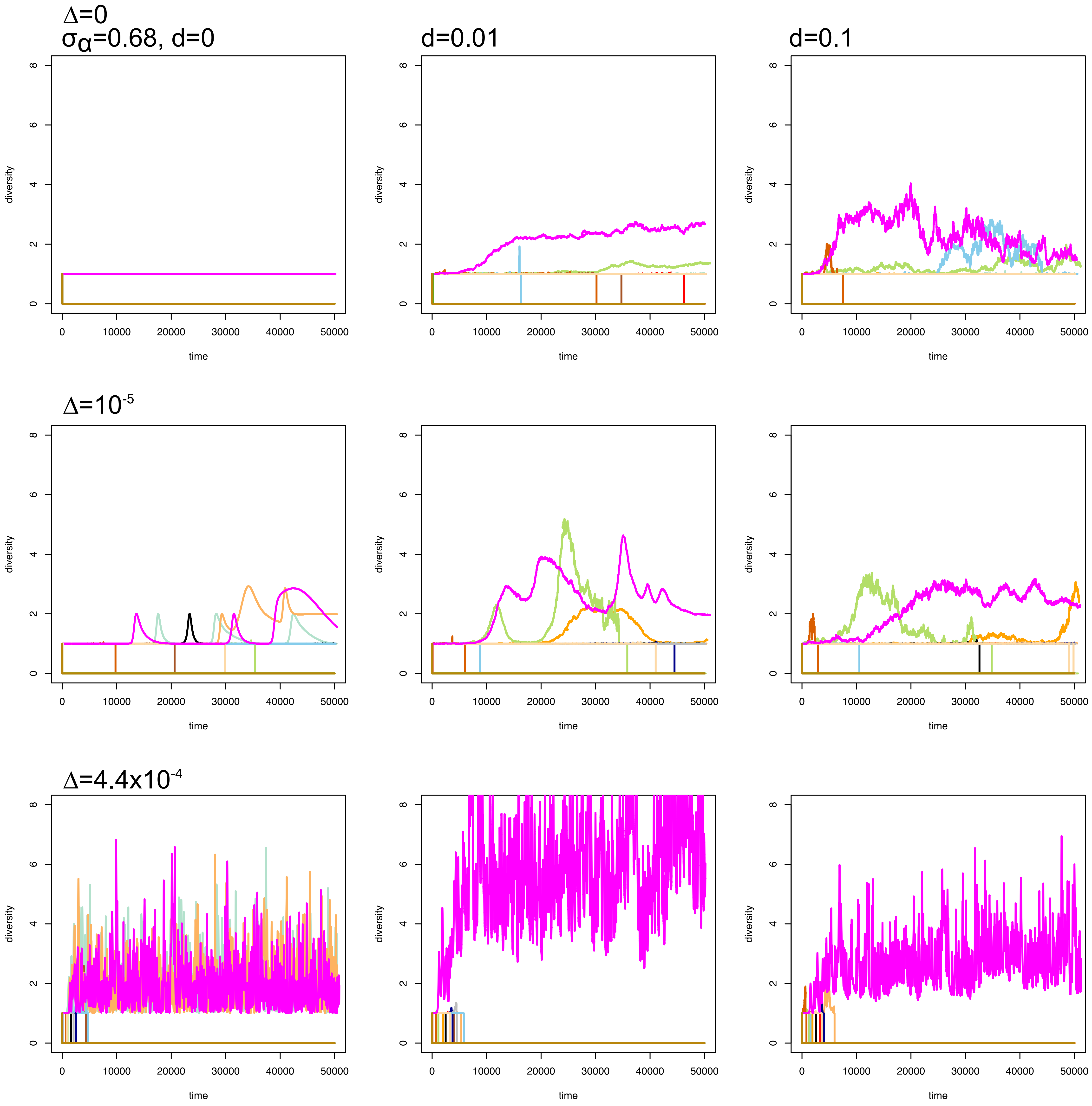

### S10

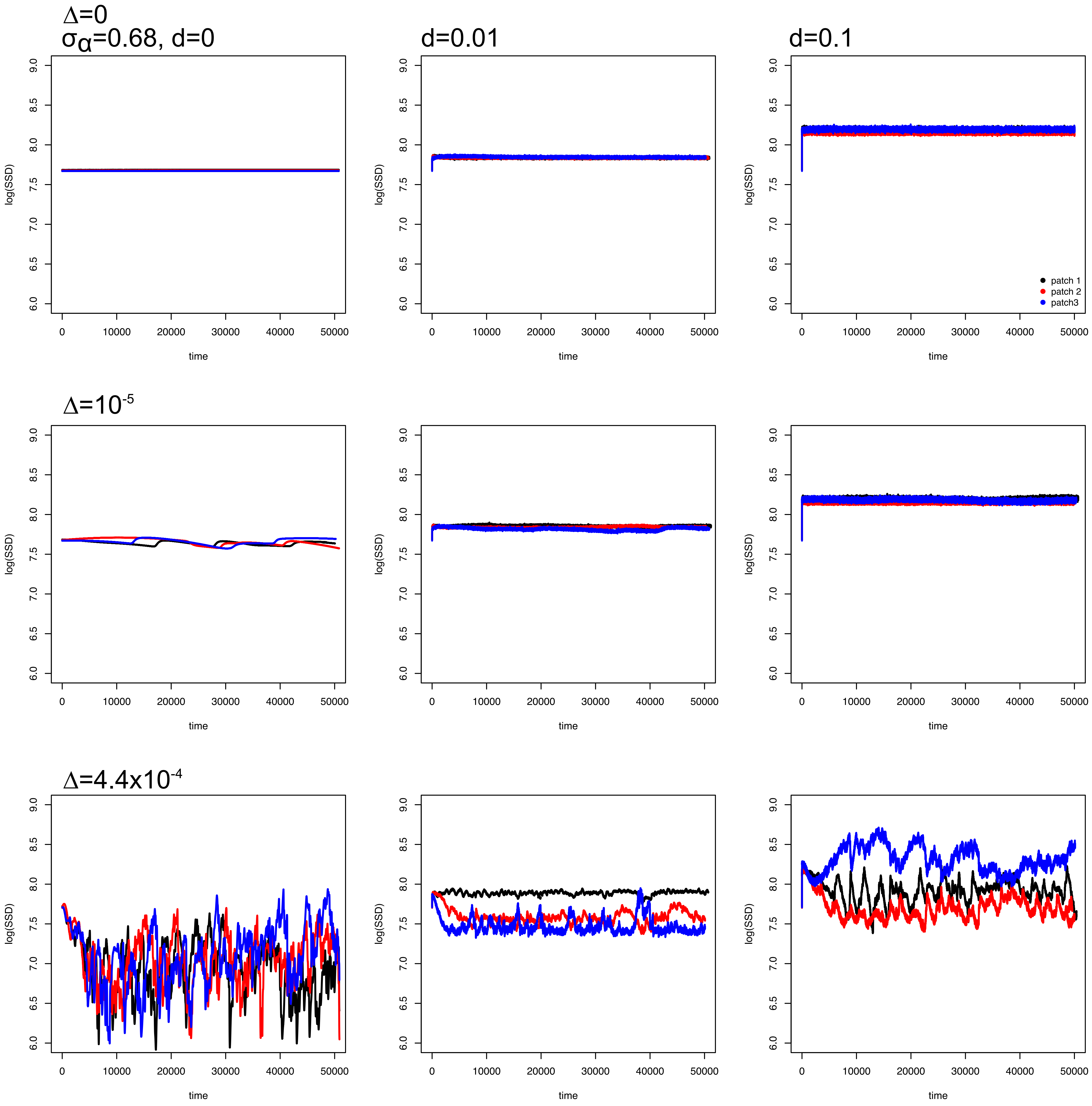

### S11

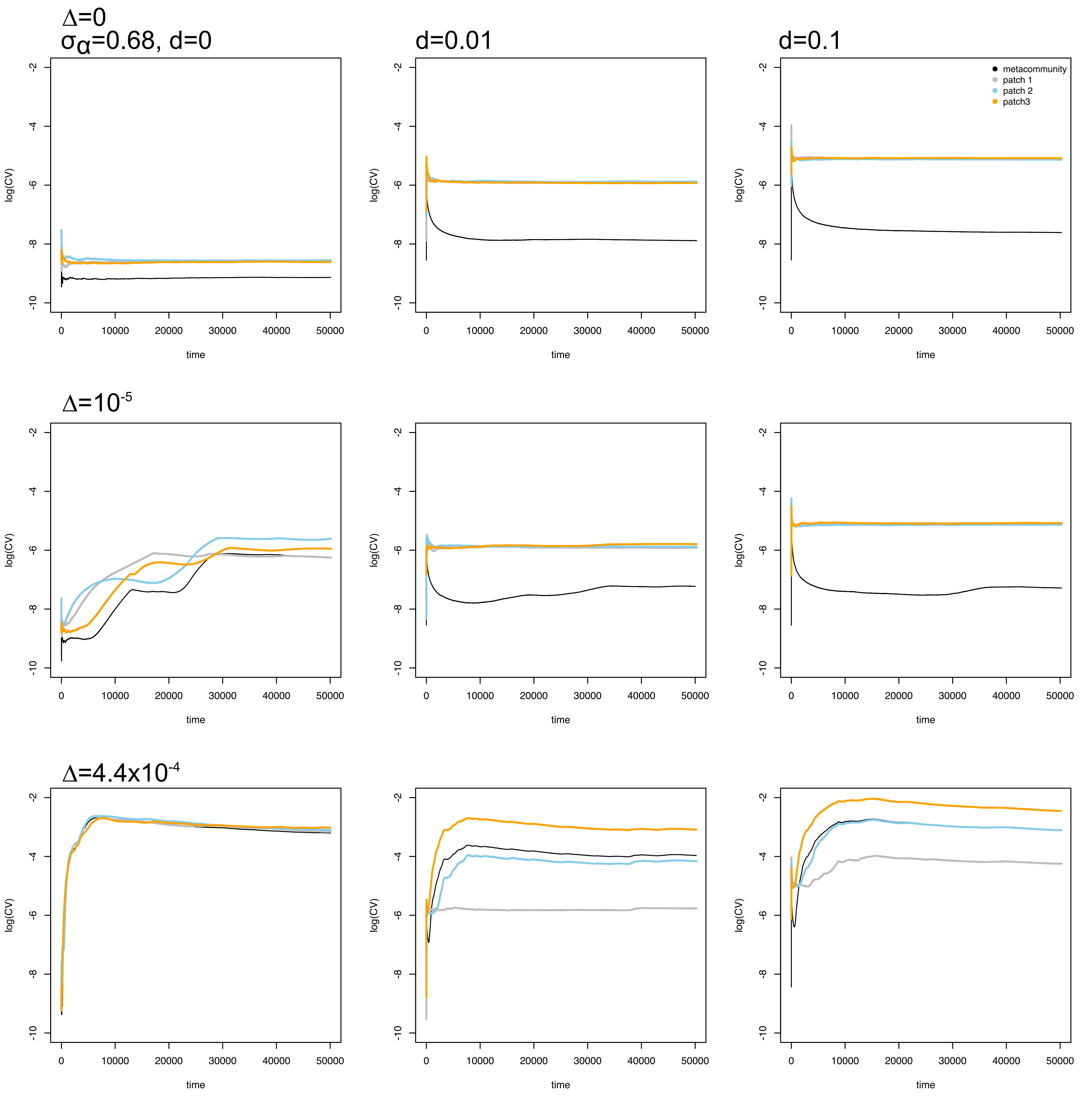

### S12

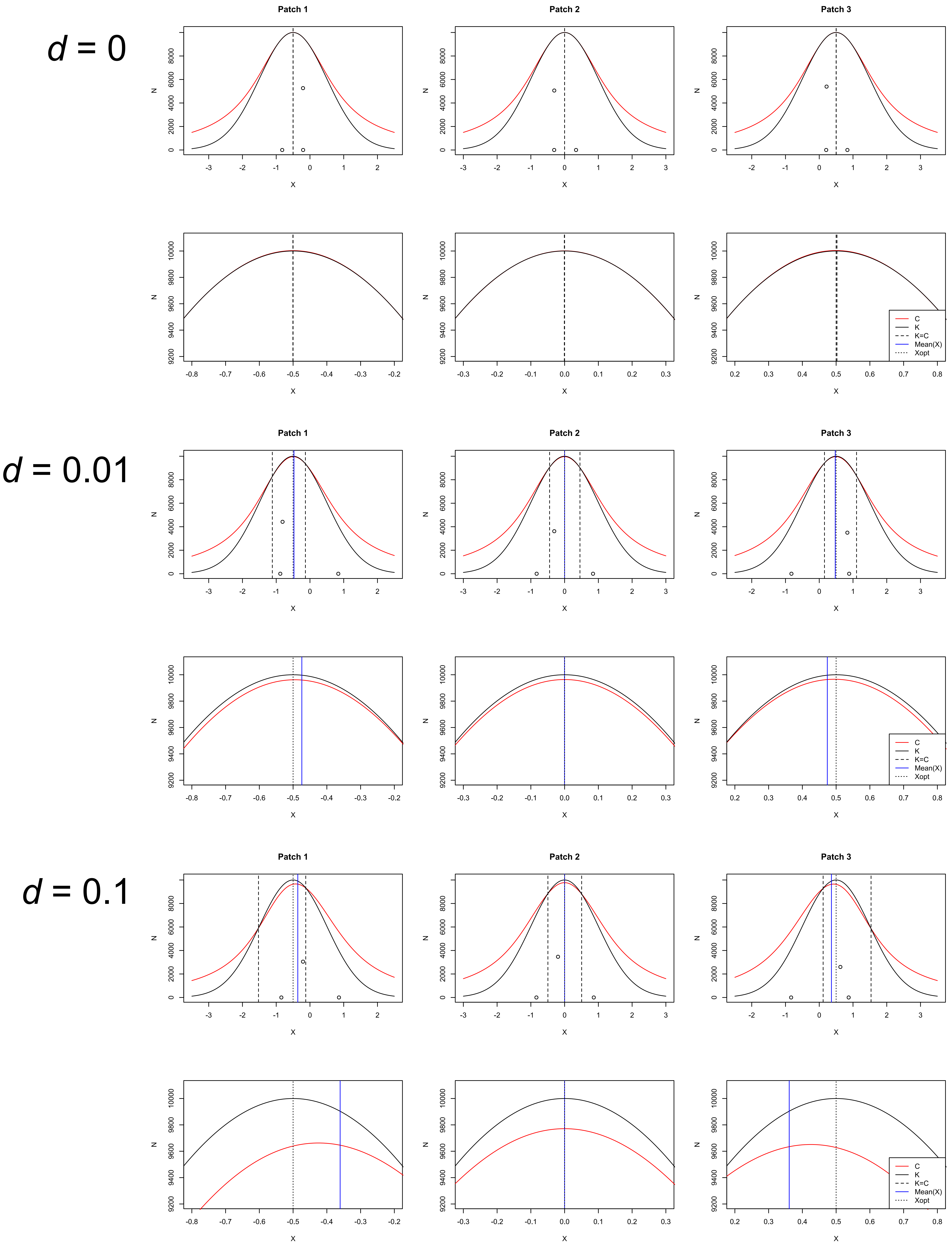

### S13

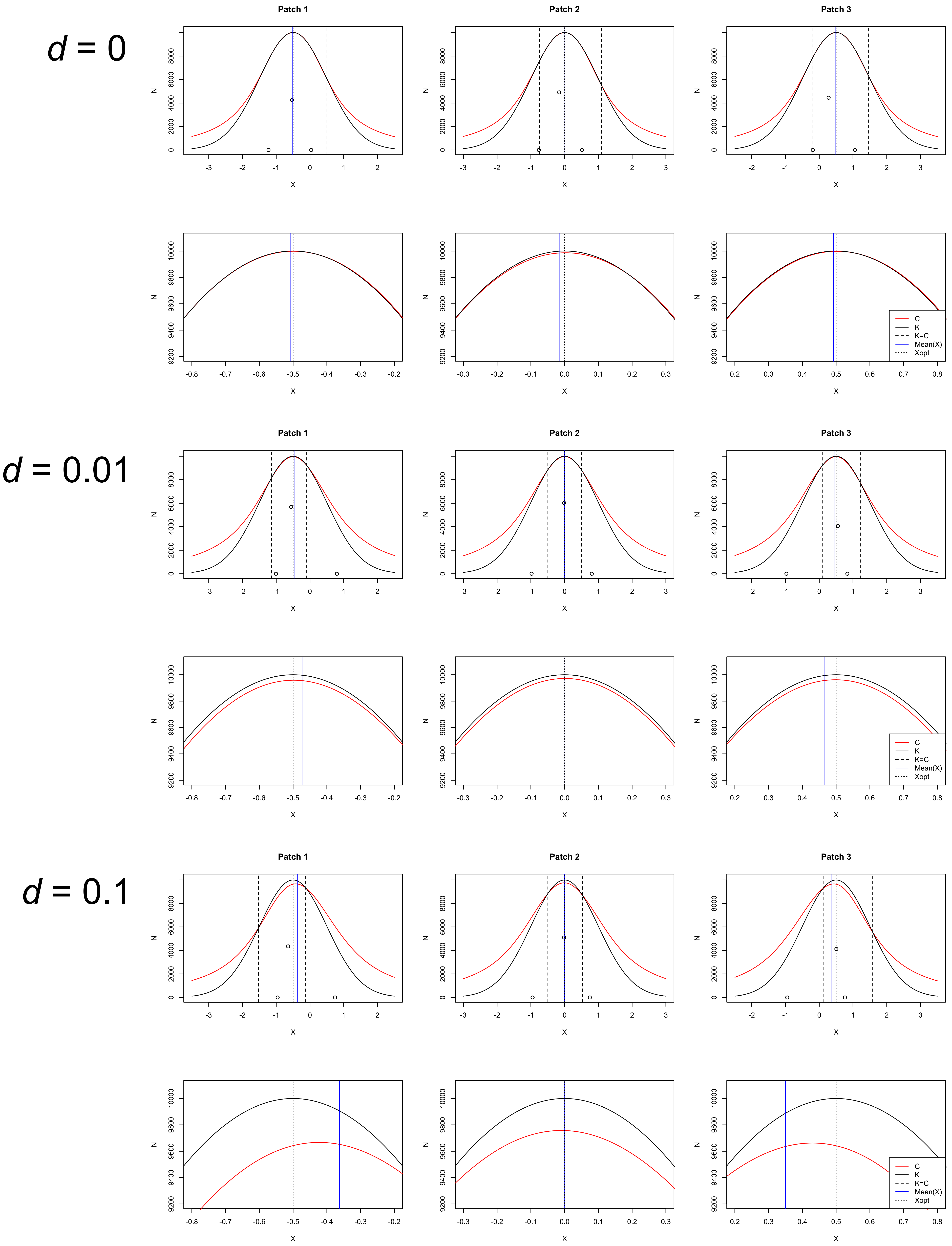

### S14

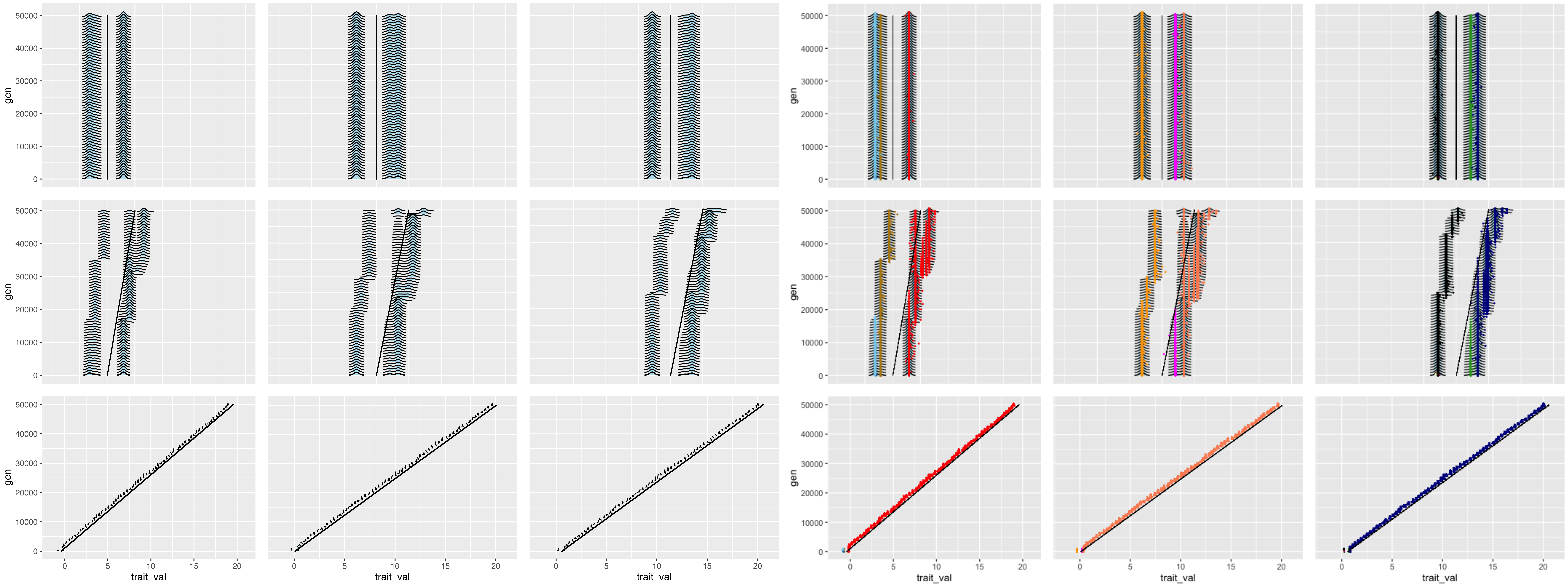

### S15

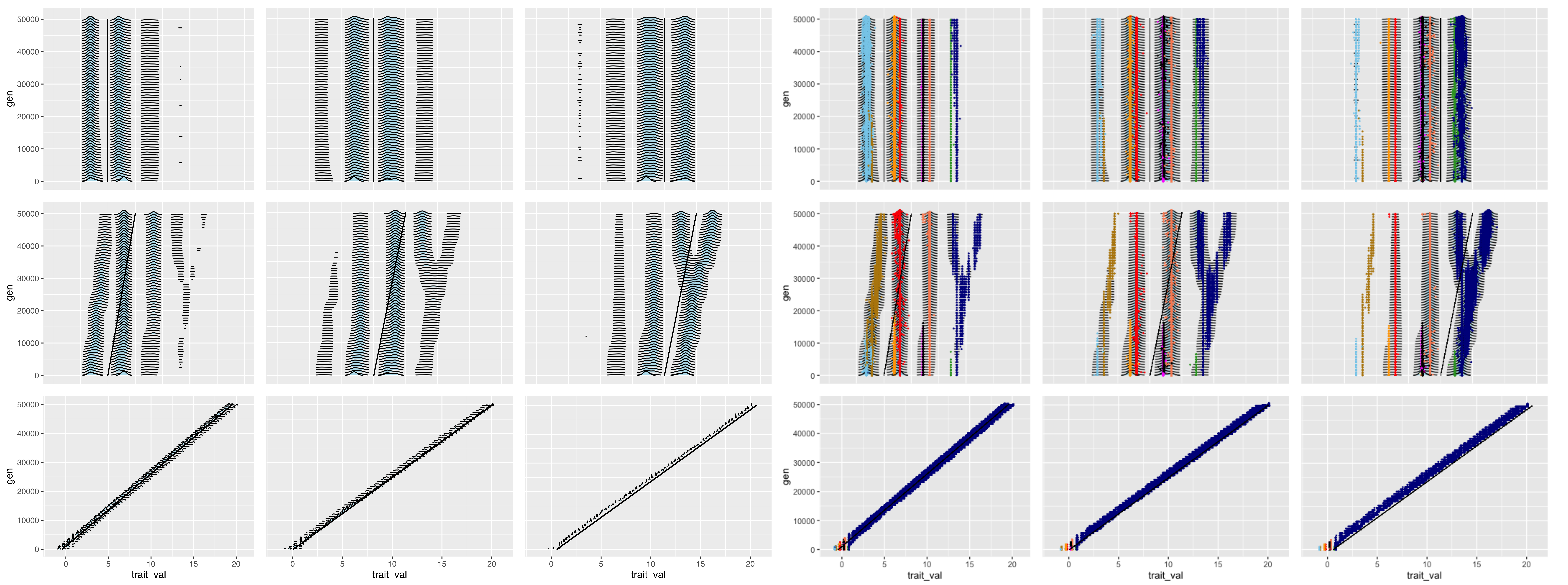

### S16

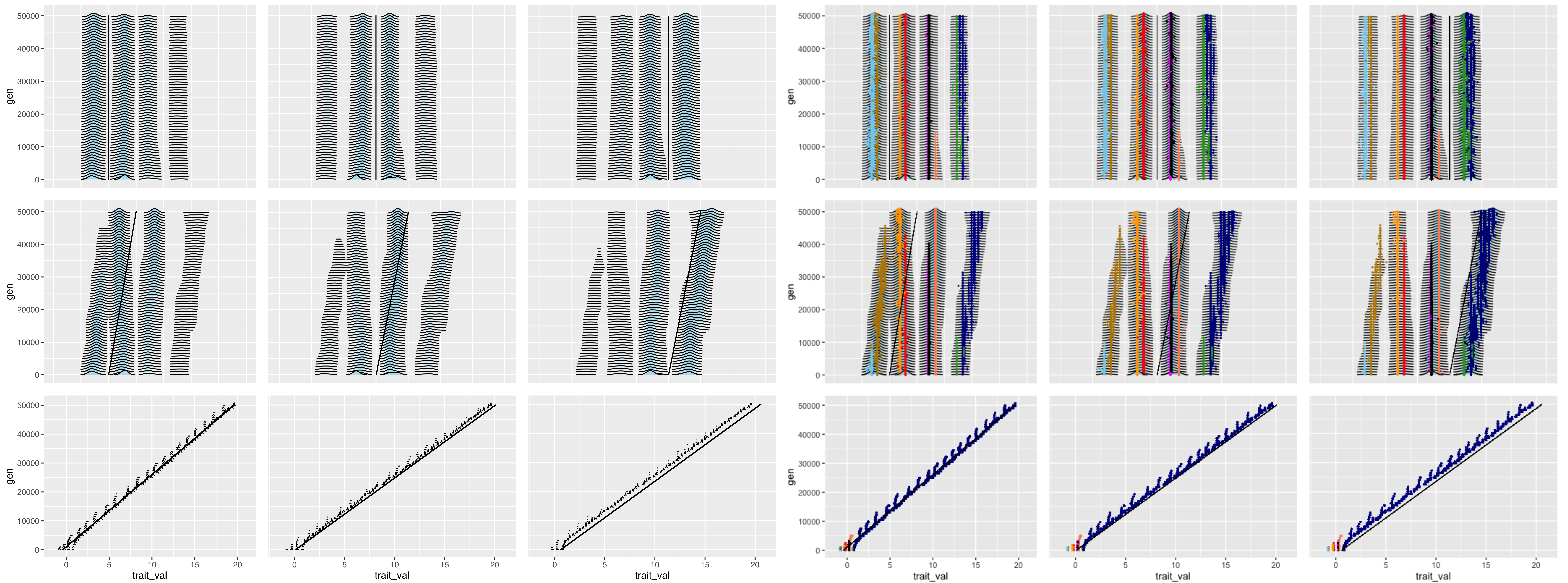

### S17

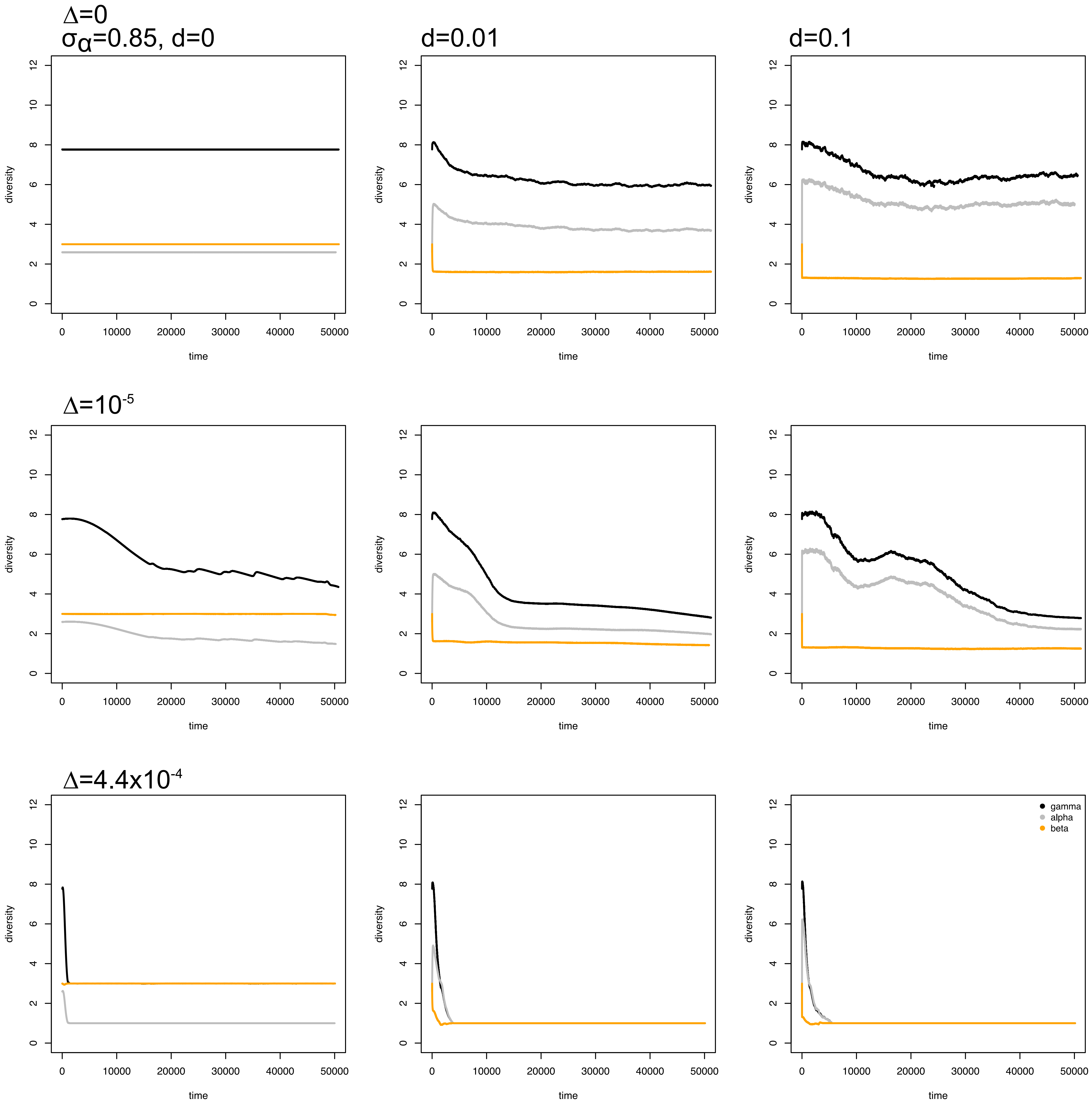

### S18

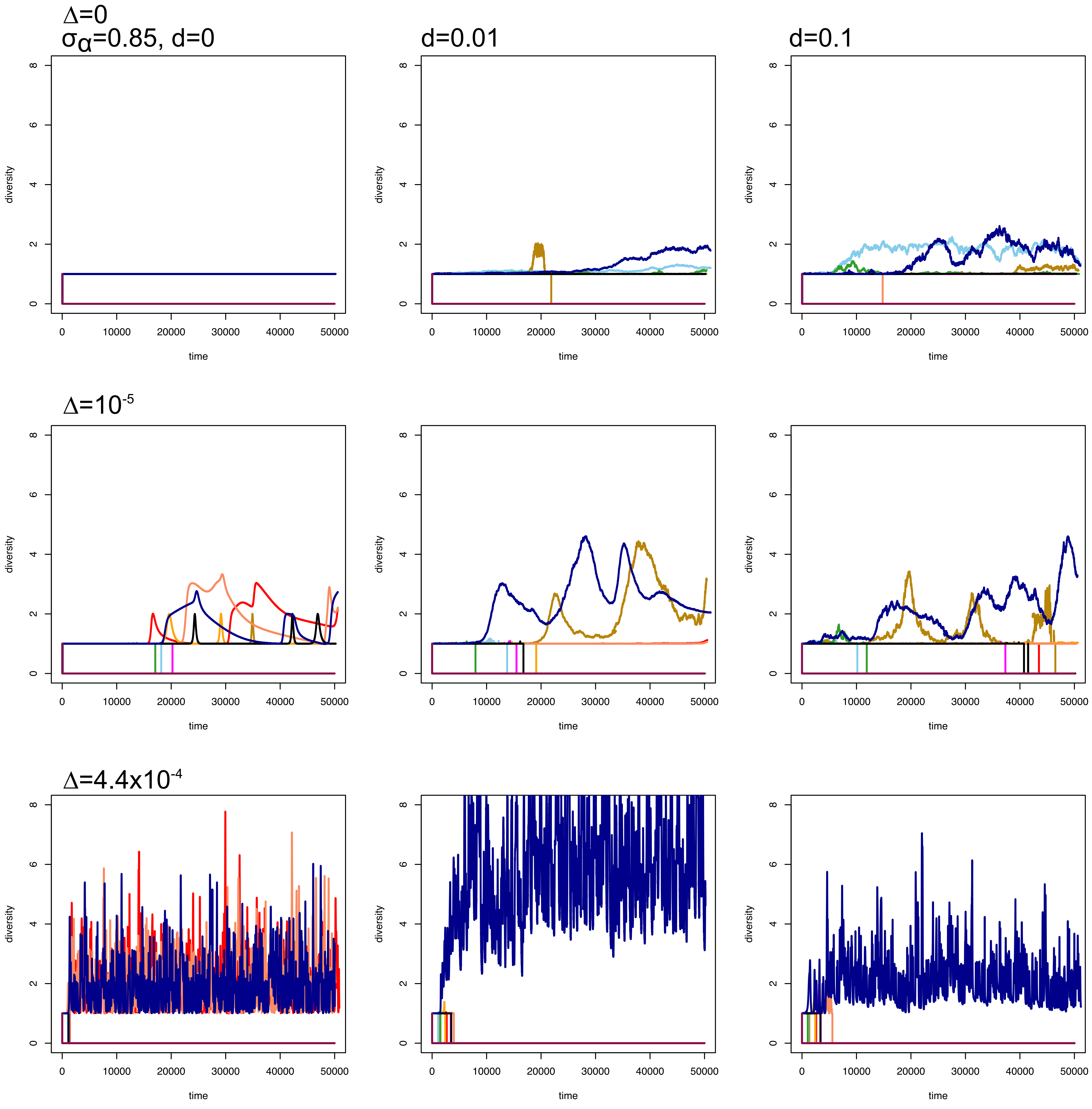

### S19

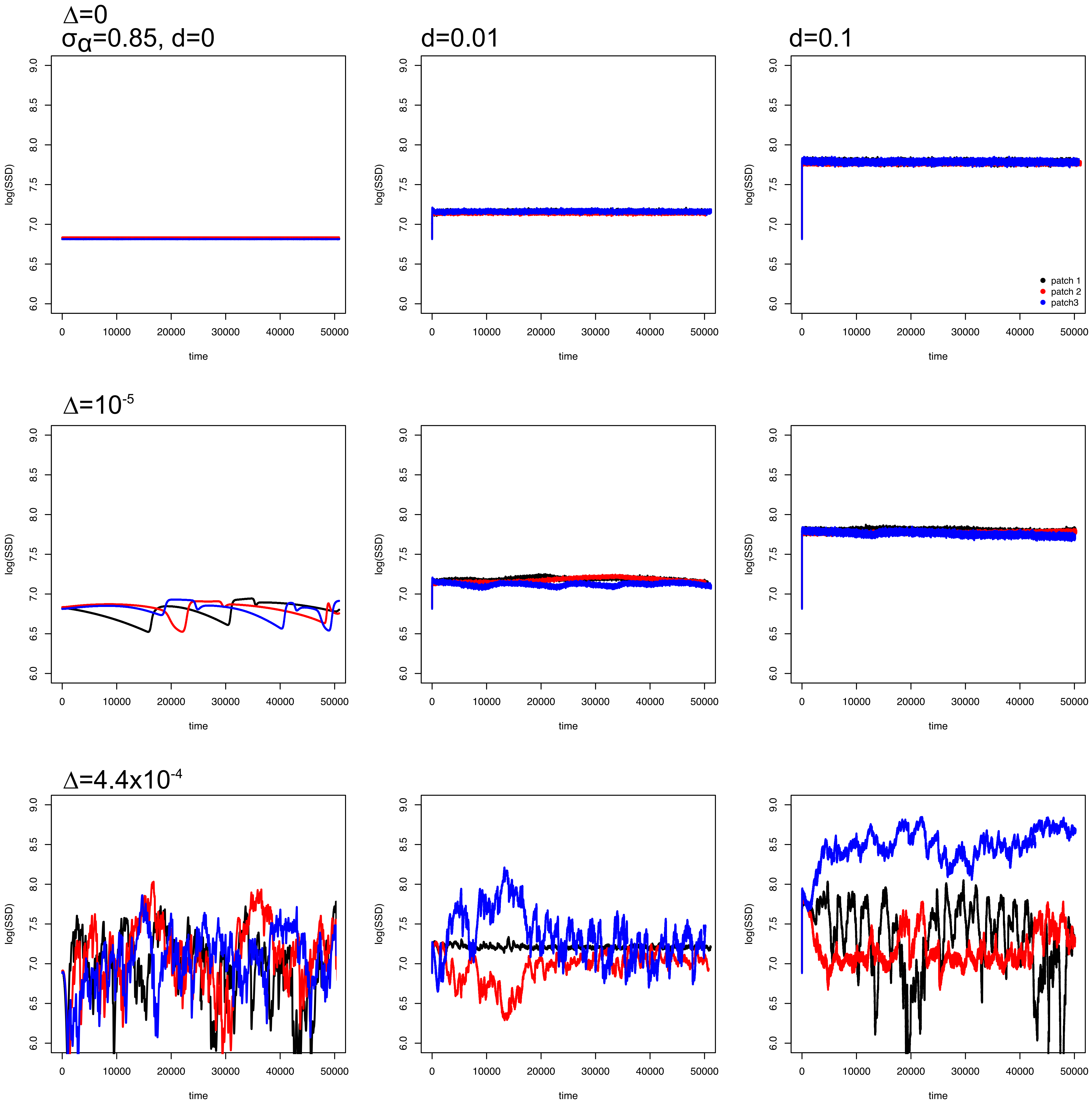

### S20

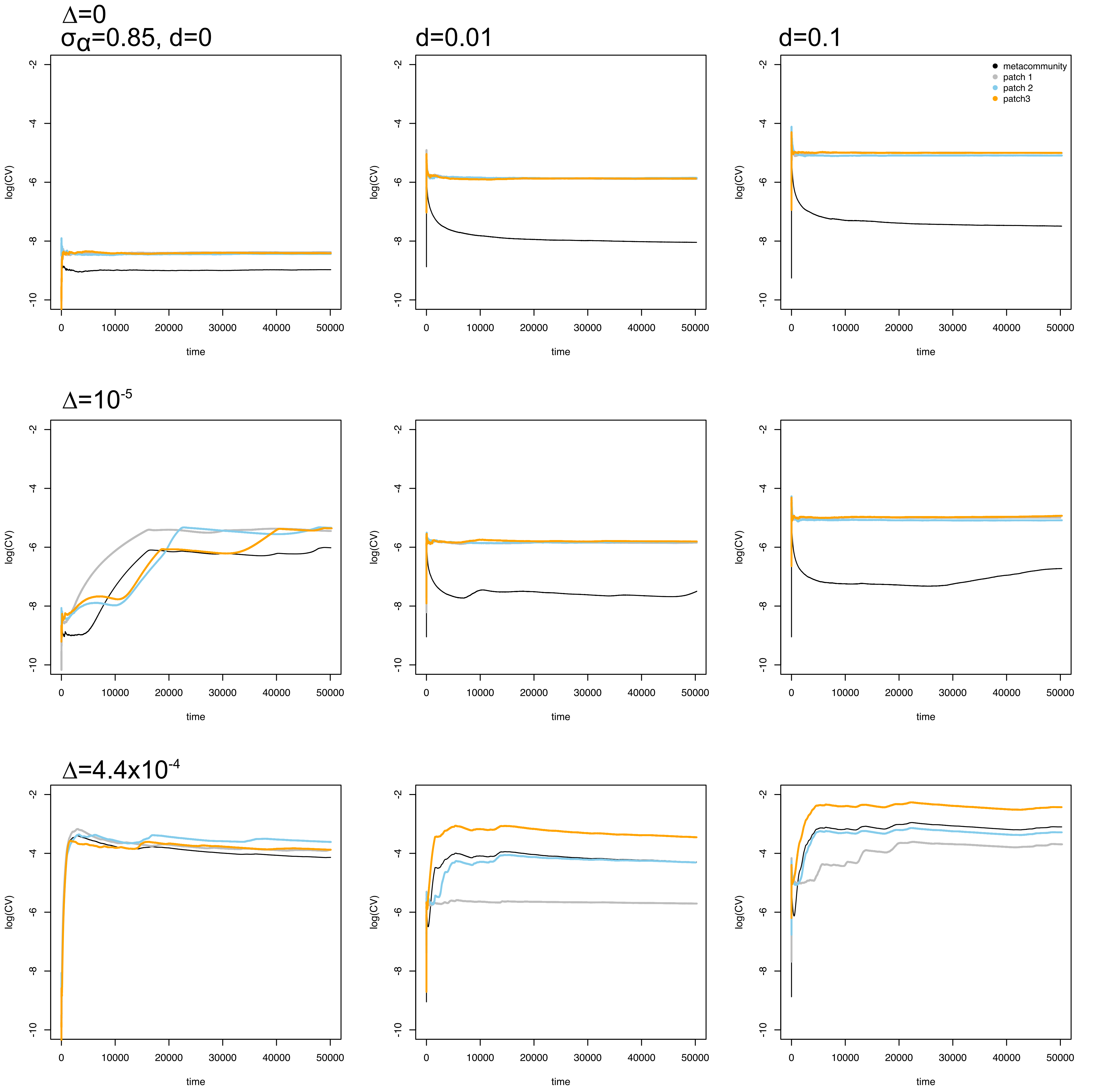

### S21

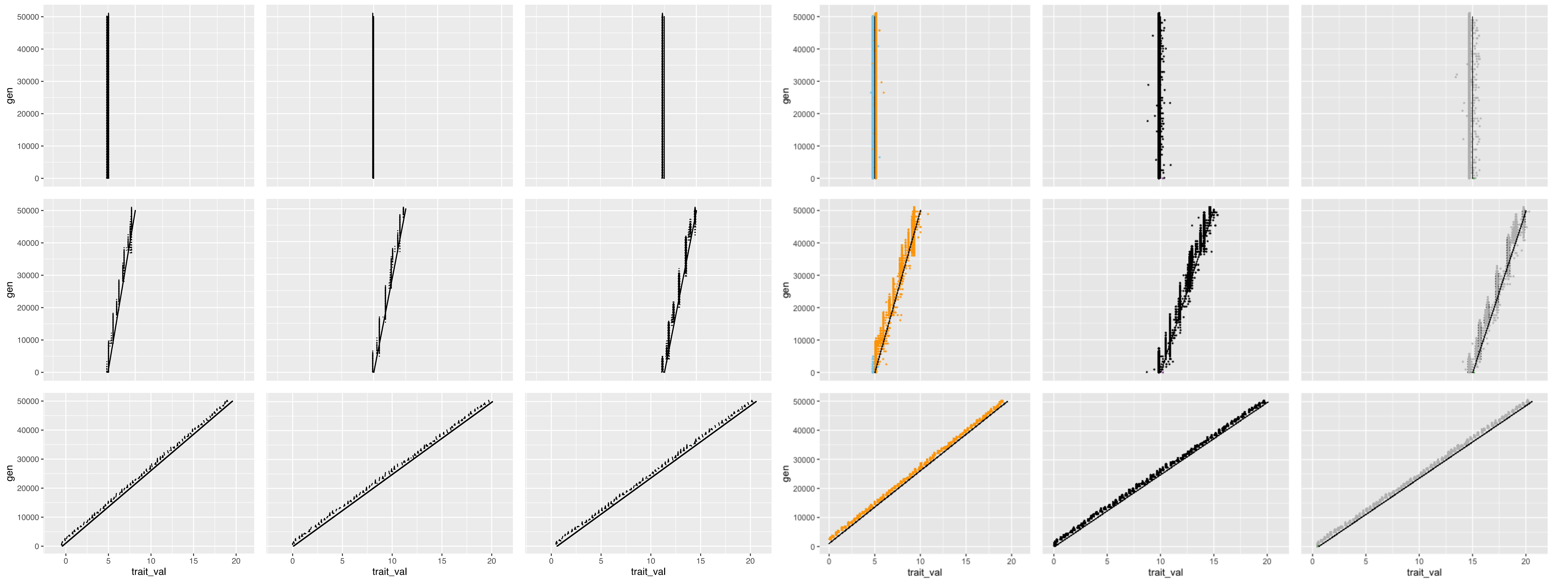

### S22

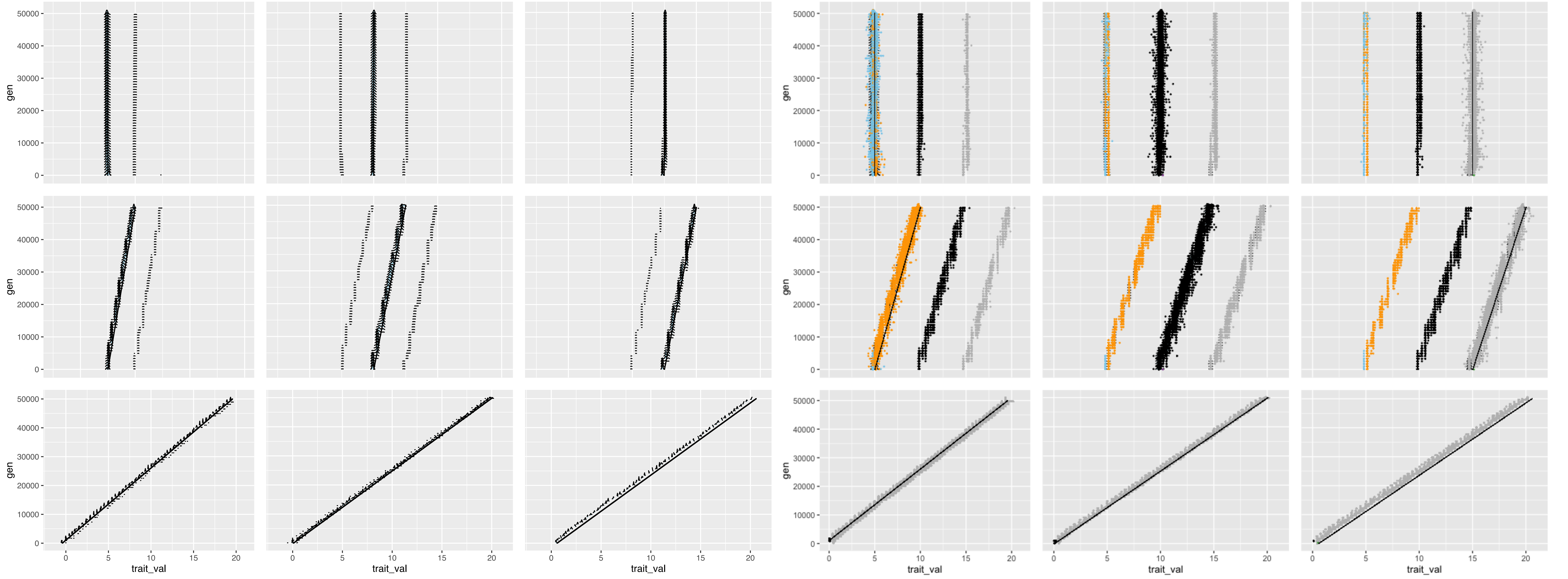

### S23

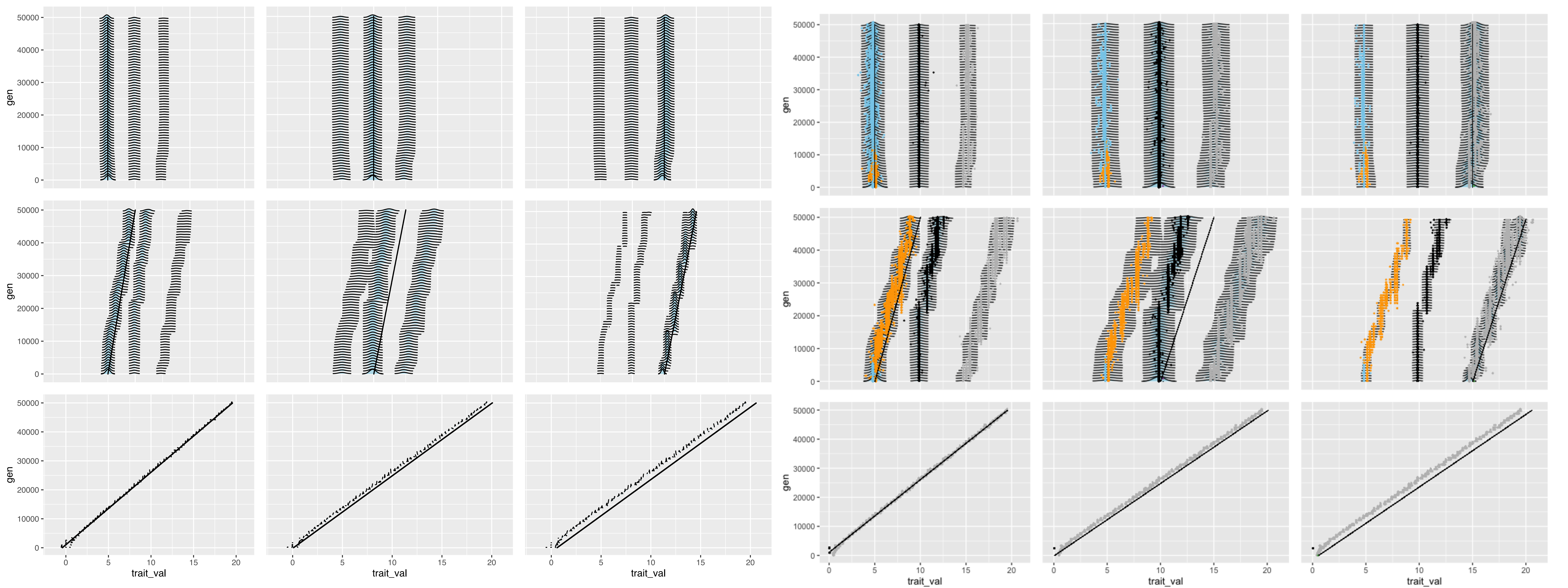

### S24

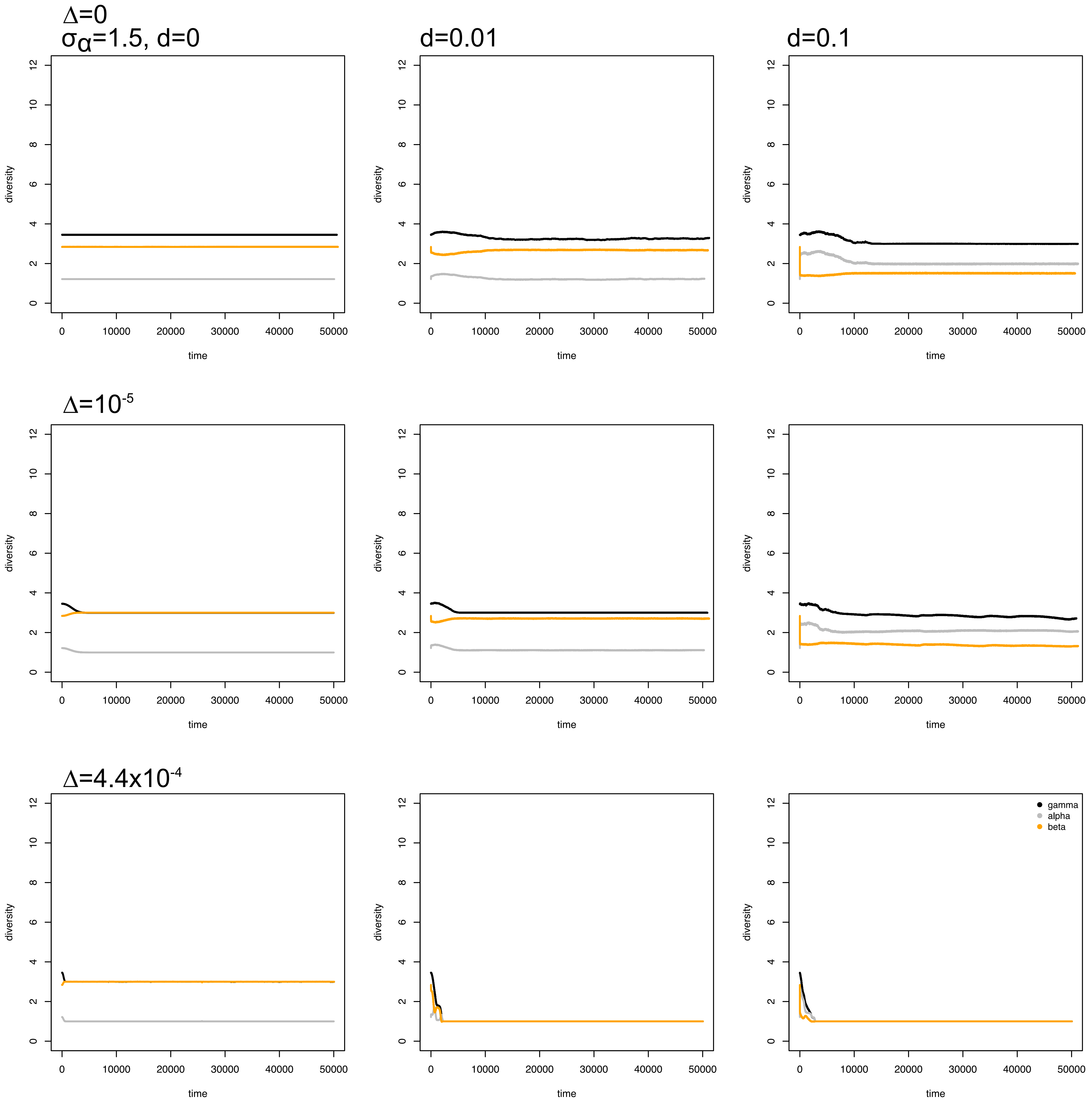

### S25

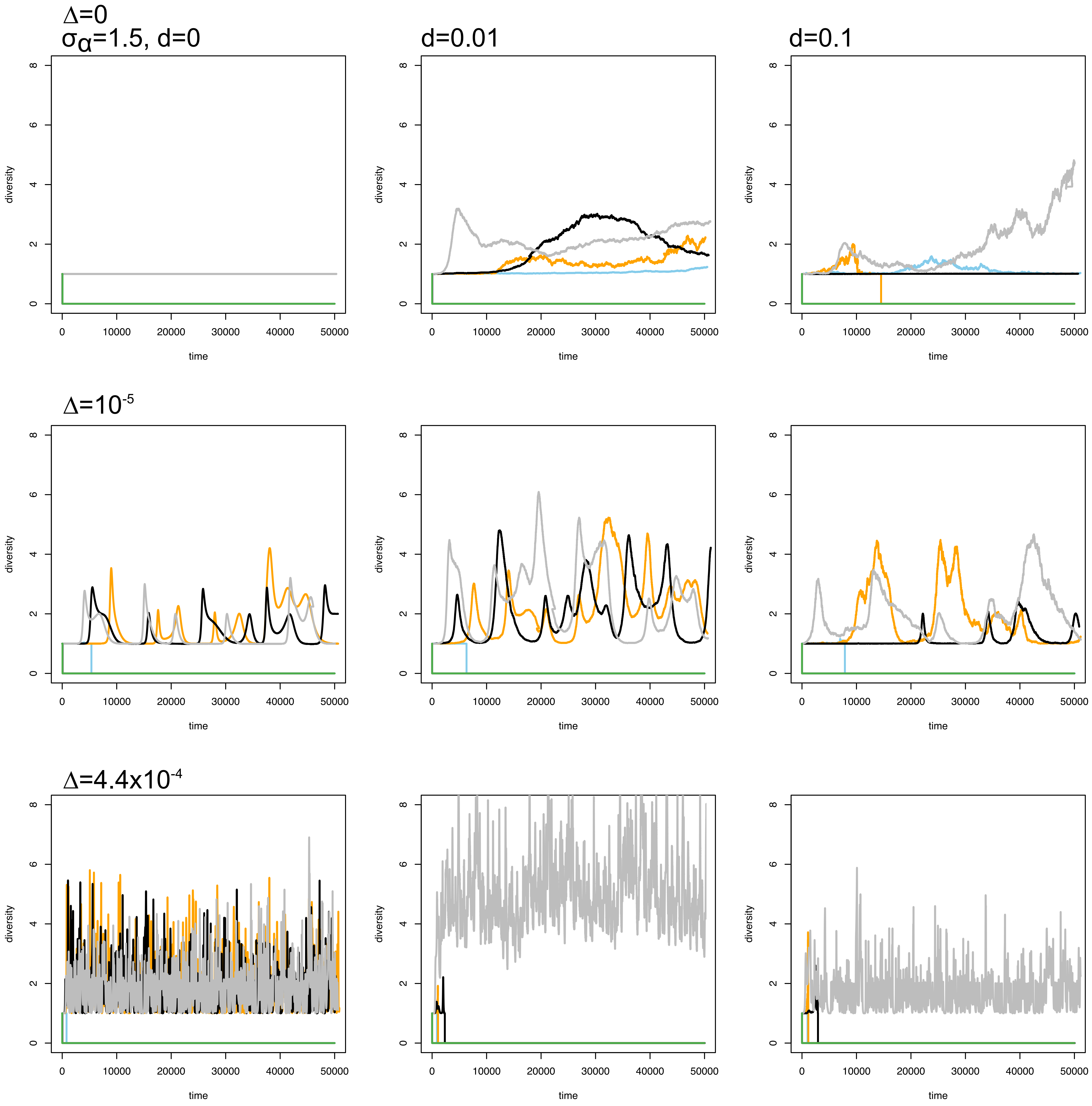

### S26

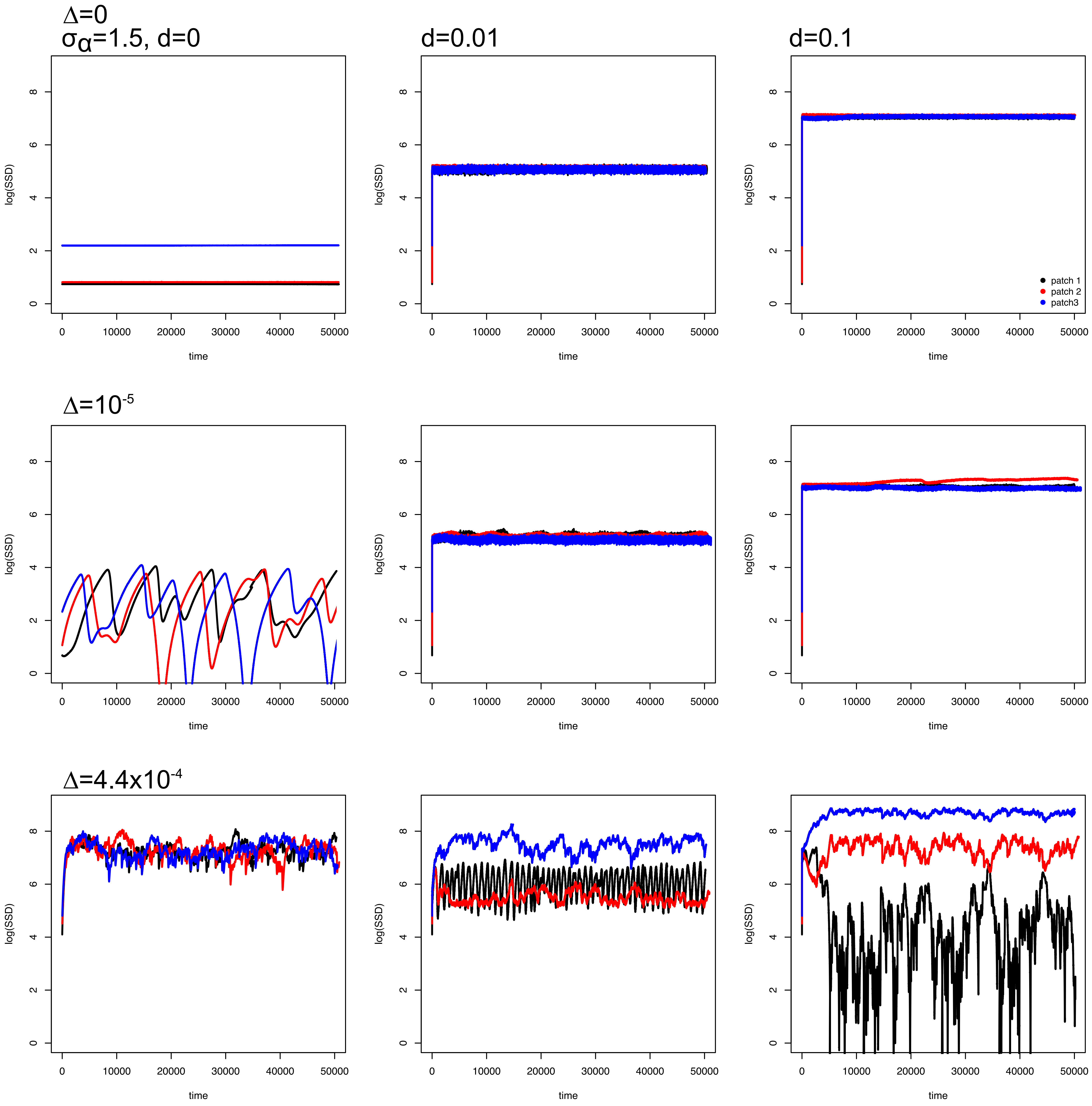

### S27

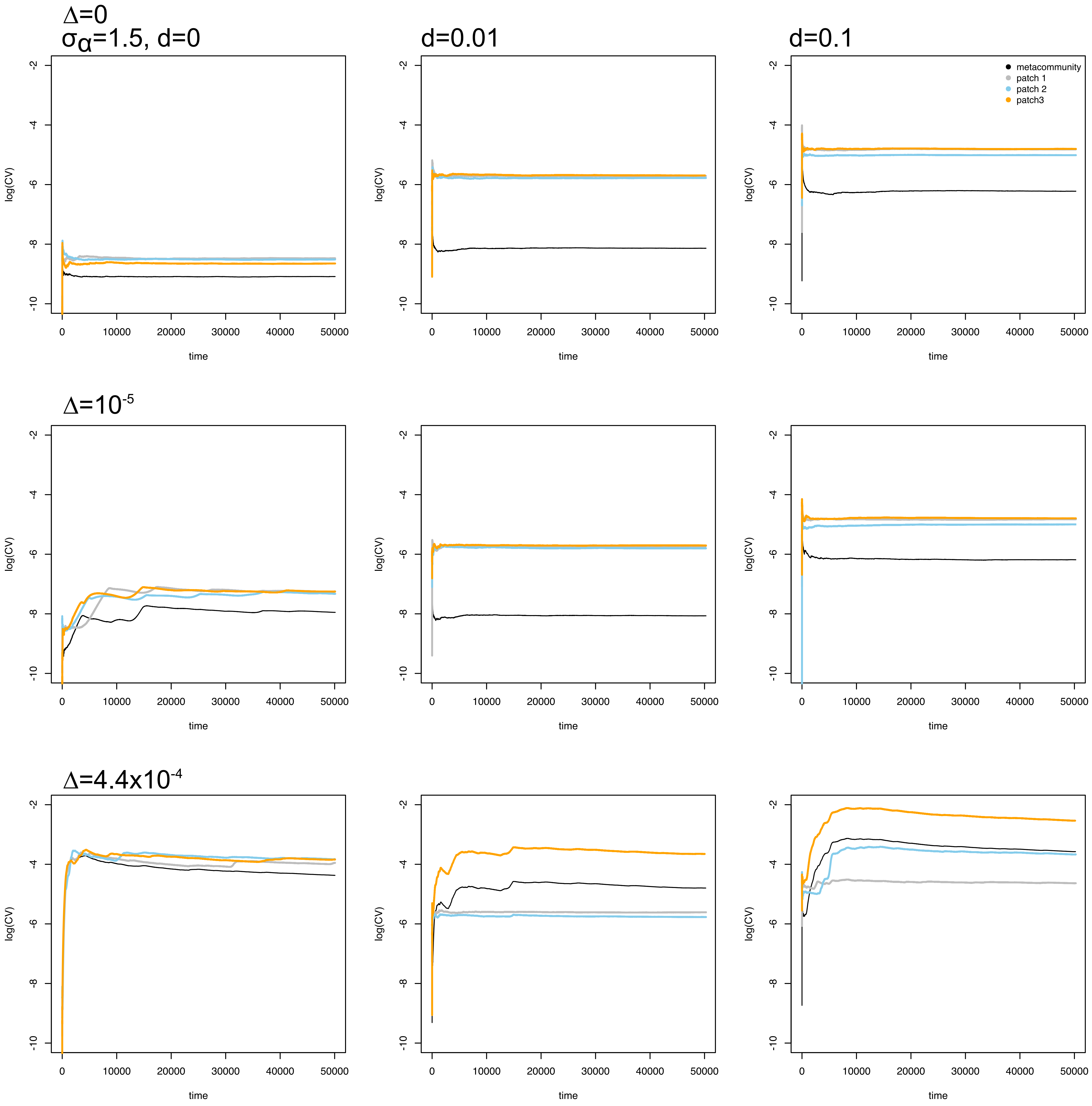
